# Hippocampal gamma oscillations form complex ensembles modulated by behavior and learning

**DOI:** 10.1101/2022.10.17.512498

**Authors:** Vincent Douchamps, Matteo di Volo, Alessandro Torcini, Demian Battaglia, Romain Goutagny

## Abstract

The hippocampus and the entorhinal cortex display a rich oscillatory activity, believed to support neural information processing in key cognitive functions^1^. In the hippocampal region CA1, a “slow gamma” rhythm (30-80 Hz) generated in CA3 would support memory retrieval whereas a “medium gamma” rhythm (60-120 Hz) generated in the entorhinal cortex would support memory encoding^2,3^. However, descriptions involving discrete gamma sub-bands can only partially account for the haphazard diversity of oscillatory behaviors observed in individual recordings during spatial navigation behavior. Here, we stress that transient gamma oscillatory episodes at any frequency or phase relative to the ongoing theta (4-12 Hz) rhythm can be recorded at any layer within CA1. Eventually, the commonly reported averages are dominated by a minority of very strong power events overshadowing gamma heterogeneity. Nevertheless, we show that such gamma diversity can be naturally explained by a simple mechanistic model, and that behavior-related information (position within a maze) can be decoded from most individual gamma events, despite their low power and erratic-like nature. Our results indicate that behavior specifically shapes ensembles of irregular hippocampal gamma oscillations, in a way which evolves with learning, depends on the hippocampal layer and is hard to reconcile with the hypothesis of rigid, narrowly tuned gamma sub-bands. Beyond randomness, the pervasive gamma diversity may thus reflect complexity at the “fringe-of-synchrony”^4^ likely functional but invisible to classic average-based analyses.

---

Coherent oscillations of neuronal activity are ubiquitous across brain spatial and temporal scales^5,6^. Oscillations at different frequencies have been associated with the formation of sensory or behavioral representations^7,8^, in the temporal organization of complex codes^9^ or in the flexible routing of information between neuronal populations^10^. The possible functional roles of oscillations have been particularly investigated in the hippocampal formation, where, in the CA1 area of the dorsal hippocampus, convergent inputs could be disambiguated by the interaction of gamma and theta oscillations: different gamma-frequency carriers, timed at different phases of the ongoing global theta oscillations, would mediate information from different sources^2,11^. Hence, slow gamma (gamma_S_; 30-80 Hz) predominates in the CA1 stratum radiatum (*rad*, where the inputs from CA3 are localized) mostly at the trough/descending phase of CA1 pyramidal layer theta. On the other hand, medium gamma (gamma_M_; 60-120 Hz) predominates in the CA1 stratum lacunosum moleculare (*l-m*, where the inputs from the entorhinal cortex layer 3 are localized), preferentially at the peak of CA1 pyramidal layer theta^2^. According to this prevalent model, layer-specific gamma oscillations in CA1 would identify the temporal dynamics of the afferent inputs, mediating specific memory-related processes (encoding for gamma_M_ *vs* retrieval for gamma_S_^11^).

Such a model, appealing for its simplicity and the link it proposes between distinct functions and discrete gamma sub-bands, may however fail to capture fully the richness of CA1 theta-gamma interactions. Recent studies investigating gamma oscillations at the theta cycle timescale reveal indeed a more dynamic and diverse landscape of gamma oscillations, with a broader variety of possible associations between gamma frequencies, theta phase and anatomical layer of occurrence (see^12^ for a recent review). Yet, these studies continued to yield a classification of hippocampal gamma into distinct sub-types, reporting a multiplicity of supposedly typical average theta-gamma patterns: from two^3^ to three^2^ or more^13,14^ gamma sub-bands. Here, we refrain from distinguishing sub-types, acknowledging that even stochastic-like oscillations with fluctuating frequency and irregular timing can self-organize to process information^4^. We therefore characterize in detail the properties of individual transient gamma events, without ignoring their broad and ubiquitous variability, which may be informative about behavior rather than merely noise.

## Gamma diversity is present in every CA1 layer

To characterize theta-gamma diversity, we analyzed local field potentials (LFPs) simultaneously recorded in the dorsal hippocampal CA1 area using 16- or 32-channel silicon probe (n=5 mice; **Fig. 1a**). For every channel, we spectrally decomposed the LFP into its main frequency components through an unsupervised algorithm (EEMD approach^15^, **Extended Data Fig. 1a**). We then computed the Current Source Density (CSD) signals using composite gamma LFPs, that is, the sum of the components peaking within a broad gamma band (30-250 Hz; Extended Data **Fig. 1b-d**). Such approach avoids any filtering within narrow gamma bands imposed *a priori*. An analogous procedure was used to construct a theta composite signal from the hippocampal fissure (4-12 Hz; **Extended Data Fig 1d**; fissure theta shows larger, more defined theta cycles than pyramidal-layer theta but with a 180° phase-shift). We then performed a time-frequency analysis to segment the gamma CSD signal into short epochs corresponding to individual theta cycles (**Fig. 1b**, left). For each segment, we characterized each of its local peaks in the gamma spectrogram (**Fig. 1b**, right) as a multidimensional vector (i.e., a *gamma element*) describing its amplitude, frequency and phase of occurrence relative to the coincident theta cycle (3 gamma features), as well as the amplitude, frequency and asymmetry of this theta cycle (3 theta features). We restricted the extraction to the four strongest gamma elements per theta cycle, obtaining thousands of gamma elements per channel and mouse (see theta and gamma counts on **Extended Data Fig 2**).

**Figure 1.**
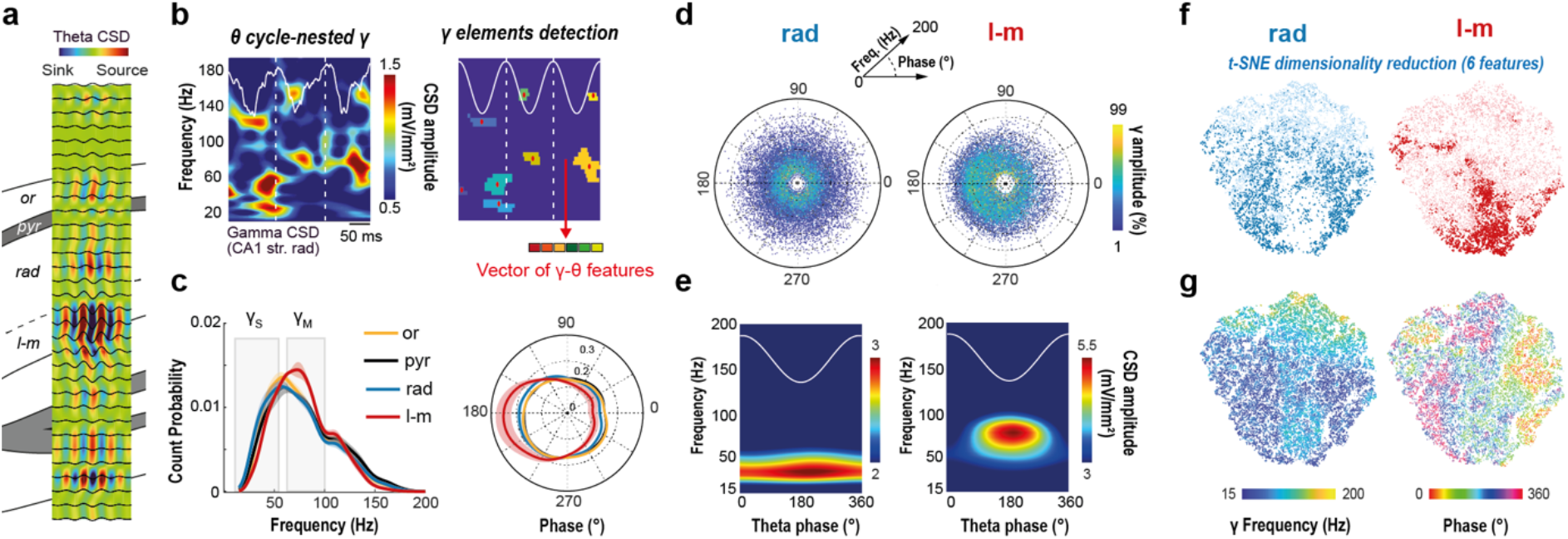
Hippocampal CA1 layers have overlapping gamma frequency and phase distributions. **a,** Electrodes along the silicon probe were localized in the different layers of the dorsal hippocampus using various indices to identify the hippocampal fissure, including the location of the maximum theta power and of the largest sink in the average theta-triggered CSD. **b,** *Left*: The gamma composite CSD wavelet spectrogram from each channel was first segmented into theta cycles (two consecutive peaks) from the theta composite signal recorded in the hippocampal fissure (white overlay). *Right*: local gamma peaks within the spectrogram were then detected within each theta cycle via a patch detection algorithm. These “gamma elements” were then characterized extracting a vector of six features: three gamma features (amplitude, frequency and theta-phase of the gamma element) and three theta features (amplitude, frequency and symmetry of the coincident theta cycle). **c**, Average distributions (mean pdf ± SEM; *n*=5 mice) of gamma elements frequency and theta phase for each CA1 layer. Even if significant differences can be found between these distributions (see Extended Data Figure 5 for details), whatever the layer considered, most of the gamma elements frequency distributions were spread across broad and overlapping ranges of frequencies, encompassing both the classical gamma_S_ and gamma_M_ sub-bands definition (full width at half maximum range: oriens (*or*), 35-97 Hz; pyramidale (*pyr*), 34-102 Hz; *rad*, 31-97 Hz; *l-m*, 38-100 Hz). **d,** A joint representation of the three gamma features emphasizes the haphazard diversity of frequency (radius) and phase (angle) between gamma elements (dots) recorded in both the *rad* and *l-m* layers, especially at low amplitude (color: percentile of gamma amplitude). **e,** Average theta-gamma spectrograms, on the contrary, put forward a marked distinction in frequency (and phase in a lesser extent) between the *rad* and *l-m* layers, suggesting these are respectively largely dominated by gamma_S_ and gamma_M_ oscillations. The apparent conflict between the representations in panels **d** and **e** is explained by the fact that average spectrograms are dominated by strong amplitude events. This is well visualized by dimensionally reduced representations (**f, g**) of the six-dimensional vectors describing gamma elements (obtained via a distancerespecting t-SNE algorithm). **f**, Gamma elements from the *rad* and *l-m* layers cover similar areas in their joint bidimensional projection. However, the elements with high gamma amplitude (top 30%, dots with darker shade) occupy complementary zones for the two layers. **g,** A color-coding by gamma frequency and phase of the same bidimensional projection shows that these strong elements tend to be: of gamma_M_ type at theta trough, for the *l-m* layer; and gamma_S_ at most phases, for the *rad* layer. These minorities of strong gamma elements are thus precisely the ones giving rise to *rad* and *l-*m average spectrogram peaks in panel (**e**). Panels **b-e**: examples from a representative mouse (mouse #3).

According to the dominant view of a theta-phase and frequency specificity of the gamma contents between hippocampal layers, we calculated the mean probability density function of these two gamma features per layer across mice (**Fig. 1c**). Unexpectedly, a substantial overlap was observed between layers for both variables, although the *l-m* presented slightly more gamma_M_ events as well as more phase-locking to the theta trough. We thus considered the joint distribution of the three gamma features for all the gamma elements per layer and animal (**Fig. 1d** for an example; **Extended Data Fig. 2** for all mice and layers), with a similar conclusion: gamma elements were broadly scattered in both frequency and theta-phase, although with a relatively stronger concentration of these in the *l-m*. Such diversity was confirmed even when extracting gamma elements with alternative techniques (e.g. filtering or independent-component analysis, **Extended Data Fig. 3a**) or from publicly available state-of-the-art recordings in rats^2^ (**Extended Data Fig. 3b**).

However, when computing the *rad* and *l-m* average theta-gamma spectrograms using the same theta cycles than for the gamma elements characterization (**Fig. 1e**), we found that they were compatible with the classic, previously reported dichotomy between a gamma_S_-dominated *rad* and a gamma_M_-dominated *l-m* (**Extended Fig. 2** for all mice and layers). In fact, our count approach revealed an increasing divergence between these layers in their frequency and theta-phase modes as the analysis was restricted to gamma elements with gradually stronger power (see details statistics on **Extended Data Fig. 4**). The discrepancy between the two approaches (“count vs average”) thus indicates that average theta-gamma spectrograms are biased by only a minority of high-power transient gamma events.

This impression was confirmed by a dimensionality reduction analysis in which we visualized in two dimensions the landscape of observed multi-dimensional gamma elements through a distance-preserving t-distributed stochastic neighbor embedding (t-SNE) nonlinear projection^16^. Strikingly, the domains covered by the projection of *rad* and *l-m* elements were largely overlapping, at the exception of the gamma elements with the highest power (see **Fig. 1f** for a representative mouse, *rad* and *l-m* are jointly projected but plot separately for visual clarity). In **Fig. 1g**, we represent as well, on the same dimensionally reduced representations, the frequencies and phases of different elements (see **Extended Fig. 5** for other features and mice), confirming once again the wide diversity of features and the lack of a simple way to discriminate between *rad* and *l-m* elements in terms of few features only. Finally, we took advantages of recent publicly available recordings made in mice undergoing transient deafferentation of CA1 via silencing of the entorhinal cortex and/or CA3 inputs^17^. Again, we confirmed that both inputs send gamma elements scattered in phase and frequency, at the notable exception of the high amplitude ones which exhibit stronger locking (see the difference maps of **Extended Data Fig. 6**).

In conclusion, we found no evidence for narrow gamma bands. The actual observations are on the contrary more compliant with a description in terms of diverse ensembles of transient gamma oscillations, widely scattered in frequency, phase and other features.

## Diverse gamma ensembles are expected at the “fringe-of-synchrony”

The observed diversity of gamma elements puts out a challenge to structure-driven views in which oscillations with different frequencies would be generated by distinct source populations^18,19^. Eventually, through a simple spiking model for generic balanced excitatory-inhibitory populations, we show that gamma diversity, rather than surprising, should be expected, as it robustly emerges for most parameter combinations provided the network remains not too far, but still below a transition to strongly synchronized oscillatory firing (i.e. at the “fringe of synchrony”, rather than in a regime with fully-developed synchrony). We considered a network with thousands of randomly interconnected excitatory (E) and inhibitory (I) quadratic integrate-and-fire (QIF) neurons^20^, driven by an external theta-modulated current input and we simulated unit activity and the associated LFP-like signals (**Fig. 2a**). **Fig. 2b-c** shows a representative raster plot of activity and the corresponding LFP spectrogram. Extracting gamma elements from simulated theta-gamma spectrograms yields a diversity of gamma frequencies and phases comparable to actual data (**Fig. 2d**). Interestingly, diversity in gamma oscillatory events is a robust property that can be obtained without the need of model parameters fine-tuning but rather arise for a very broad range of conductance and drive intensity. Gamma frequency diversity, quantified in terms of spectral entropy, remains uniformly high over most of the represented parameter space, colocalizing with regimes of weakly synchronized and low-amplitude oscillations (triangle, circle and square example working points in **Fig. 2e** and **Extended Data Fig. 7a**). Spectral entropy drops uniquely above a transition to a strongly synchronized oscillatory regime, characterized by higher amplitude fluctuations of the mean membrane potential (star symbol in **Fig. 2e**). Wherever spectral entropy is high, oscillations are transient and display scattering in frequency and phase like in empirical data (four paradigmatic cases are characterized in **Extended Data Fig. 7c**). Yet, the relative probability of occurrence of elements with higher or lower frequencies can be smoothly controlled, by varying the strength of the coupling of excitatory to inhibitory neurons (triangle circle, and square symbols in **Extended Data Fig. 7b-c**). In sum, a local balanced E/I circuit at the fringe-of-synchrony is not expected to generate a gamma rhythm with a narrowly tuned frequency but, on the contrary, a diverse ensemble of transient gamma events with dynamically adjustable average frequency.

**Figure 2.**
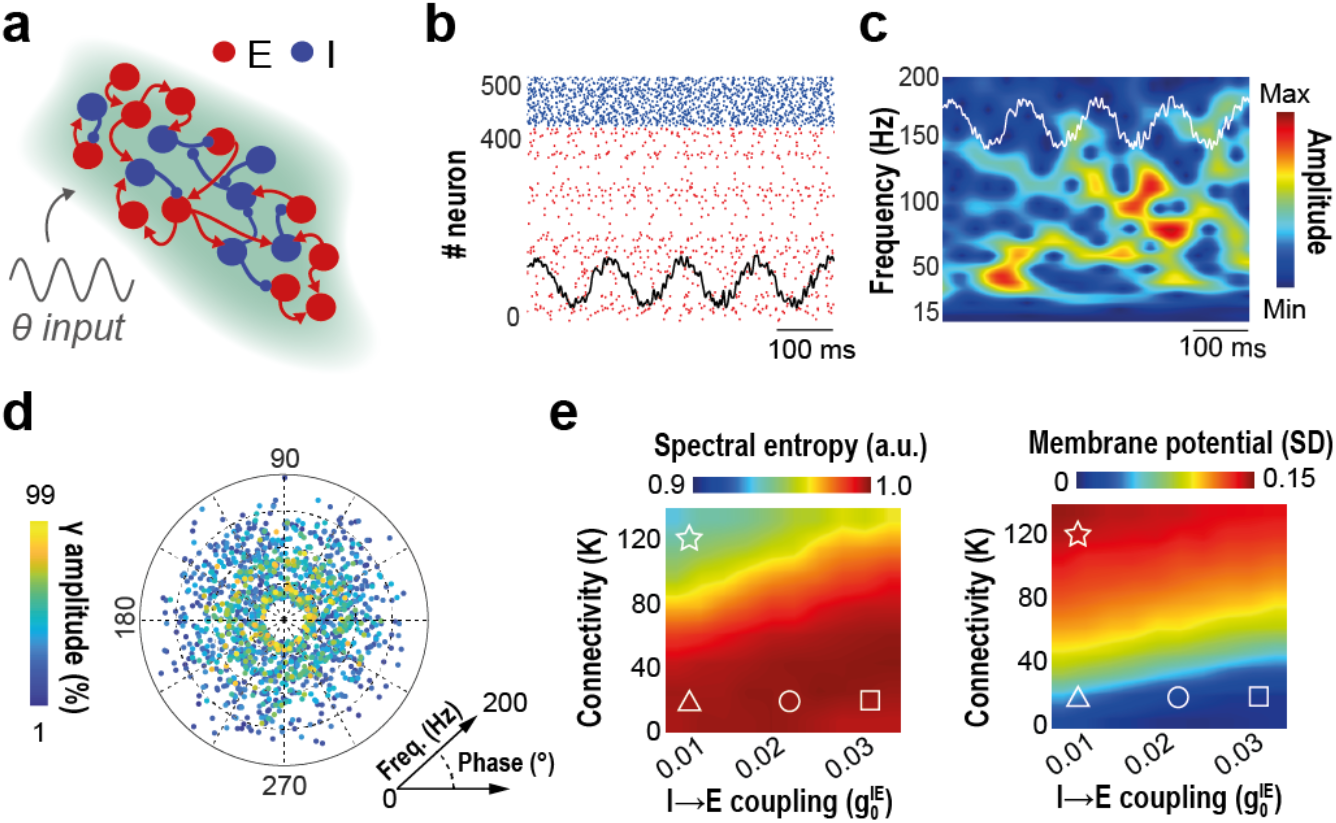
Large diversity of gamma elements is expected at the fringe-of-synchrony. **a,** We generated simulated LFP-like signals using a computational model of a generic local circuit, generating gamma oscillations and driven by an external theta-modulated input current. The model network included thousands of randomly interconnected spiking excitatory (*E*) and inhibitory (*I*) neurons. **b,** Typical raster plot of the spiking activity of selected neurons, with superposed trace of the associated LFP-like signal computed from the model. **c**, Spectrograms of the gamma composite component of simulated LFP-like signals reveal the existence of transient gamma oscillatory events at variable frequencies and phases (cf. Figure 1b). **d,** Gamma elements extracted from simulations have a landscape of feature diversity comparable to real recordings (individual dots are individual gamma elements, same representation as in Figure 1d). **e,** The diversity and frequency distribution of simulated gamma elements depend on model parameters, such as the density of within-population connectivity *K* and the average strength of *I*-to-*E* synaptic coupling. The parameter-dependency surface of spectral entropy (left; larger when power is spread more uniformly across frequencies) show that narrower-band oscillations with a more precise frequency occur only when connectivity *K* is very large (star working point). However, in this case, the level of synchronization in the population would be unrealistically large, as manifested by a very large signal standard deviation (right; larger when population oscillations have stronger amplitude). On the contrary, the simulations of panels (**b-d**) have been realized at the circle working point, for which activity is more irregular and only sparsely synchronized (fringe-of-synchrony regime, robustly leading to higher spectral entropy and lower signal standard deviation). Changing the strength of *I*-to-*E* synaptic coupling maintains the system within a fringe-of-synchrony regime but shifts the mode of the generated gamma elements’ frequency distribution (cf. Extended Data Figure 7; triangle working point: mode in gamma_S_ range; square working point: mode in gamma_M_ range).

## Navigation behavior can be decoded from individual gamma elements

Diversity of gamma elements is thus pervasive, both *in vivo* and *in silico*. But is this diversity potentially functional? In our computational model, the diversity of gamma elements is the byproduct of balanced activity within a circuit with irregular connectivity. It is possible however that, in empirical recordings, the stochastic-like variations across theta cycles of gamma power, frequency, and phase are modulated by actual behavior. To answer this question, we trained the mice to learn a novel spatial reference memory task (**Fig. 3a**). In this task, mice seek for an appetitive target located within an 8-arm radial maze. They need to learn a unique, stable goal location over multiple days of training (10 days; 4 daily trials). The change of departure arm in each of the few daily trials enforces the comparison of allocentric cues with internal representations. We therefore attempted decoding the current position of the mouse within the maze during navigation behavior, based on the features of the simultaneously recorded gamma elements. To do so, we trained machine learning classifiers (ensembles of randomized decision trees, **Fig. 3b**) to predict the rough location of the mouse (four non-overlapping maze sections: target arm approach, target arm reward field (Reward RF), other arms’ reward fields (other RF, no reward) and rest of the maze; **Fig. 3b**) based on the six-dimensional vector parametrization of a coincident individual gamma element recorded on a specific channel. The training set for each classifier was restricted to elements from a subset of randomly chosen theta cycles, the unused elements being allotted for later cross-validation of the performance. Decoding yielded performances well above chance-level, particularly for the target arm and the reward field, for any layer within CA1 (**Fig 3c** for a representative mouse; **Extended Data Fig. 8a-e** for all mice). Given the limited modulation of the performance by the anatomical position (channel), we summarized the achieved channel-averaged performances of decoding for all mice. We showed that performance for decoding target arm, reward field and other arm end-fields were well above chance level for every mouse (**Extended Data Figure 9a**, see also the confusion matrix in **Extended Data Figure 9b**). Even if location could be better decoded from events whose amplitude belonged to the largest quartile of the amplitude distributions –the one dominating spectrograms (cf. **Fig. 1e**)–, decoding was possible even from elements at weaker amplitudes in the other quartiles, with the exclusion of only the lowest quartile of amplitudes (**Fig. 3d**, left, for target arm and reward field decoding performance by gamma amplitude quartile; **Extended Data Fig. 9c** for other locations). We then analyzed whether the performance of location decoding depends on the speed of movement of the mouse (**Fig. 3d**, right, and **Extended Data Fig. 9d**). Decodability of reward field and other arms ending zones was higher when speed was low (lowest quartile) and when speed was high (highest quartile) for the target arm. When speed was large, decoding was significant for all four considered maze locations and confusion between locations was reduced (**Extended Data Fig. 9d**). Yet, we were able to significantly decode location from elements in other speed quartiles (down to the second quartile for target and up to the fourth quartile for reward) indicating that speed is not the unique determinant of gamma element modulations by maze location (note that speed distributions over the different maze sections are not identical but, still, largely overlapping, cf. **Extended Data Fig. 9l**). Analogously, decodability was maintained across several quartiles of the other features in the gamma element parametrization, that is, gamma frequency and phase relative to theta, and theta cycle amplitude, frequency and asymmetry (**Extended Data Fig. 9e-i**). In short, decodability was not limited to narrow categories of elements with specific feature combinations, but was on the contrary rather pervasive, extending notably to gamma elements strongly deviating from spectrogram averages. In some cases, more information could even be extracted from weak than from strong amplitude gamma elements (as when decoding presence in the reward-less other RF locations, **Extended Data Fig. 9c**).

**Figure 3.**
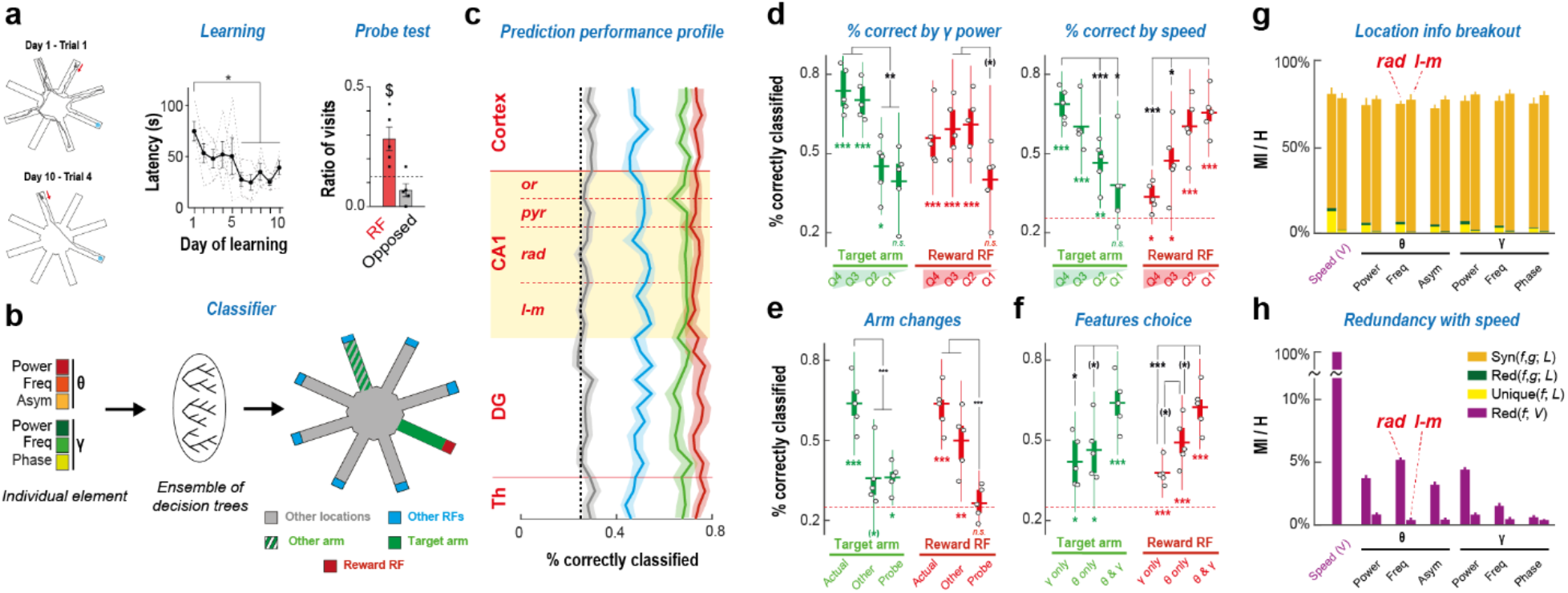
Location during exploration behavior can be decoded from individual gamma elements. **a,** In our spatial navigation task, reward is located at the end-box of a fixed target arm in a radial maze. The mouse enters the maze from a different arm at every trial. Few trials are performed every day, over several days. Left: example trajectories at different days of learning. Middle: the latency to reward decreases across days (ANOVA, p < 0.02), indicating that mice learn the task. Right: In probe trials, no reward is given at the reward location. Mice spend a larger amount of time exploring the former rewarded arm than an opposite one, denoting memory of the reward location (one-tailed t-test, p < 0.03). **b,** We trained tree-ensemble classifiers to decode rough location within the maze (target arm, reward RF, other RFs and the remaining locations) from individual gamma elements (three theta and three gamma features, cf. Fig. 1b). We also trained alternative classifiers to detect an alternative arm, remote from reward. **c,** Fraction of correctly classified locations, by maze location (colors as in **b**), for a representative mouse (mouse #3; see Extended Data Figure 8 for other mice and prediction performance with alternative feature sets and in probe trials). Different classifiers were trained for different depths along the dorsal hippocampal axis (cf. Figure 1a). Solid lines indicate average performance across all trials (shading, 95% bootstrap c.i.). Performance in detecting target arm, reward and other arms RFs was significantly above chance level (dashed black line) for every anatomical layer. **d**-**f,** Decoding performance was robust and affected by various factors (dots, performance averaged over trials and electrodes for different mice; boxes, IQRs and sample mean; whiskers, 95% sample c.i.; *, p < 0.05; **, p < 0.01; ***, p < 0.001 after Bonferroni correction; symbols in brackets denote significance only before Bonferroni correction; one-tailed t-tests of sample vs chance level; two-tailed t-tests between samples). **d,** Left: dependence on gamma amplitude. Performance is significantly higher for amplitudes in larger than smaller distribution quartiles (p < 0.002 for target; p < 0.033 for reward), however it remains significant for all but the lowest quartile (p < 0.006 for target Q3). Right: dependence on motion speed. Decoding performance for target arm (reward RF) was higher for larger (lower) quartiles of speed (p < 0.004 for target, p < 0.001 for reward), but was significant even for low (high) speed quartiles (target Q3, p < 0.0025; reward Q1, p < 0.011). **e**, In probe trials, decoding performance dropped for both target arm (p < 0.0005) and reward RF (p < 0.0001). Decodability of a generic other arm was lower than for target arm (p < 0.0027), but performance did not drop significantly for reward. **f,** When training classifiers to decode maze location based on reduced input feature sets (only gamma- or theta-related features) decoding performance dropped (e.g., when comparing gamma-only with combined theta and gamma inputs, p < 0.0199 for target arm and p < 0.0014 for reward RF). **g,** Fractions of maze location information conveyed by pair of features were large (bars, averages over mice and feature pairs, grouped by pairs including a specific feature, for representative *rad* and *l-m* channels; whiskers, 95% bootstrap c.i). Mutual information was mostly due to synergy between features, which conveyed little unique or redundant information about location**. h,** Speed accounted for a small fraction only of the variability of individual gamma element features, as revealed by normalized mutual information with speed (averages over all mice and features, for representative channels; whiskers, 95% bootstrap c.i).

To further verify that our classifier extracted genuine behavior-related information from individual gamma elements, we first modified our classifier design by training the classifier not to specially identify the target arm but a randomly chosen arm among the behaviorally non-saliant (i.e., neither departure nor target arm) ones. The decoding performance that could be reached for these generic arms was not as high as when decoding the approach to the actual target arm (although the decoding of the reward zone was not significantly changed, **Fig. 3e**, and **Extended Data Fig. 9j** *other* boxes). This indicates that classifiers can detect signatures in gamma elements –akin to an “eureka” signal– which specifically reflect behaviors observed when approaching the target arm but no other arms of the maze. Second, at the end of task learning, we performed a probe trial in which the reward was removed. Such probe condition modified the behavior during target arm approach and reward field exploration (cf. **Fig. 3a**, right), as reward was unexpectedly missing at the previously learned location and context was thus altered. As shown by **Fig. 3e** and **Extended Data Fig. 9j** (*“probe”* boxes), classifiers trained to decode target arm and reward fields in learning trials could still significantly decode transit in the target arm zone (although with a lower performance) but the performance in decoding the reward field dropped at chance level. Such pattern of performance modification was consistently observed across all recording channels and mice (cf. **Extended Data Fig. 8f-j**). Thus, behavior induced by the probe condition translates into modified gamma element signatures, since the same classifiers that decoded relevant maze locations in preceding trials could not identify them anymore in the probe trial.

Therefore, the features of individual gamma elements –very diverse, especially when gamma amplitude is low– are modulated by maze location and behavior in complex but consistent ways that machine learning classifiers can successfully identify.

## Different features of theta and gamma oscillations synergistically reflect behavior

Which features give the largest contribution to the successful decoding of maze location from diverse gamma elements? To address this question, we constructed machine learning classifiers using, as input alternative, smaller subsets of features: only the three gamma features (*gamma-only*) or only the three theta features (*theta-only*). Target arm and reward field could still be decoded above chance level based on the gamma-only or the theta-only subset of features, however, the performance dropped with respect to the original classifier, indicating that theta and gamma-related dimensions of the gamma elements convey non-redundant information (**Fig. 3f** and **Extended Data Fig. 9k**). Information theory and, specifically, the framework of Partial Information Decomposition (PID^21^) can be used to further investigate the nature of this nonredundancy. Indeed, two input features *f* and *g* (e.g., theta and gamma amplitudes) can convey: *unique* mutual information about an output feature *L* (here, maze location), conveyed by one but not the other input; *redundant* mutual information, shared by both inputs; and, beyond that, *synergistic* mutual information which the two inputs convey in joint coordination, but not when considered independently. Given the growing intricacy of the PID framework for larger groups of input variables, we computed here partial decompositions for the information about location conveyed by pairs of gamma element features.

We first averaged these PID analyses over pairs of input features pooled by feature (i.e., all pairs including gamma frequency, all pairs including gamma amplitude, etc.; see **Fig. 3g** for a synoptic view and **Extended Data Figure 10** for non-averaged pairs). The considered gamma and theta feature pairs carried on average over 75% of the information needed to perfectly specify maze location at any time, thus explaining why decoding of location is feasible. Next, we fractionated the total mutual information into the parts constituted by the unique, redundant and synergistic fractions (respectively in yellow, green and orange colors, **Fig. 3g**). Remarkably, the synergistic fraction was by far the most important, accounting in most cases for over 70% of the total information conveyed by the feature pairs about maze location. These indicate that individual theta- and gamma-related oscillatory aspects have individually complex and changing relations with maze location (hence the low unique information fractions) but that their joint patterns of general covariation do depend on it (hence the large synergistic fractions).

We also considered mutual information between maze location and speed of movement, as the distributions of the speed of movement were not completely identical for different maze sections (**Extended Data Fig. 9l**). As shown by the leftmost bar of **Fig. 3g**, predictor pairs including speed among the input variables did not convey significantly more total information about maze location than any other pair of gamma element features. The only specificity of speed as an input feature was that it conveyed more unique information about maze location than any other oscillatory feature taken individually, at least for *rad* (but not for *l-m*) gamma elements (cf. also, in more detail, **Extended Data Fig. 10**). Yet, even if feature pairs involving speed do not carry more maze location information, the encoding of maze location by oscillatory features may still indirectly reflect relations with speed, via the dependence of the oscillatory features themselves on speed. Therefore, we also computed the redundancy of speed with the other oscillatory features. Individual oscillatory features of the *rad* shared more information with speed than oscillatory features of the *l-m* (see **Extended Data Figure 10**). However, the shared information with speed never explained more than 5% of their variation entropy (**Fig. 3h**). Together these results indicate that variations of gamma element features are not completely explained by speed, but synergistically convey genuine maze location information, beyond mere speed variations across locations. The dramatic drop in inter-feature synergies observed during probe trials may thus explain the lower maze location decoding performance in these with respect to learning trials (**Fig. 3e** and **Extended Data Figure 9j** and **10**).

## Complex gamma ensembles evolve with task learning

Behavior can be decoded out of gamma elements, but is the decoding grammar similar across learning? And is the decoding performance improving with training? We explored this by training classifiers over gamma elements from trials within restricted ranges, starting from early trials and then sliding the inclusion range to the latest trials. Gamma element outstanding diversity was present at any trial range, noticeably never losing their continuous and broadly dispersed frequency and theta-phase distributions despite slight changes (**Fig. 4a**, see also the t-SNE projections in **Extended Data Fig. 6c** and polar plots in **Extended Data Fig. 11**). Maze location information could be significantly decoded from these diverse gamma elements at any trial range, although the detailed profiles of variation across learning were heterogeneous for different mice, possibly reflecting idiosyncratic navigation learning strategy. Yet, the cross-validated fraction of correct predictions was larger for late than for early trials, with a performance improvement on average of ~7% for the *l-m* and of ~15% for the *rad* layers (**Fig. 4b**).

**Figure 4.**
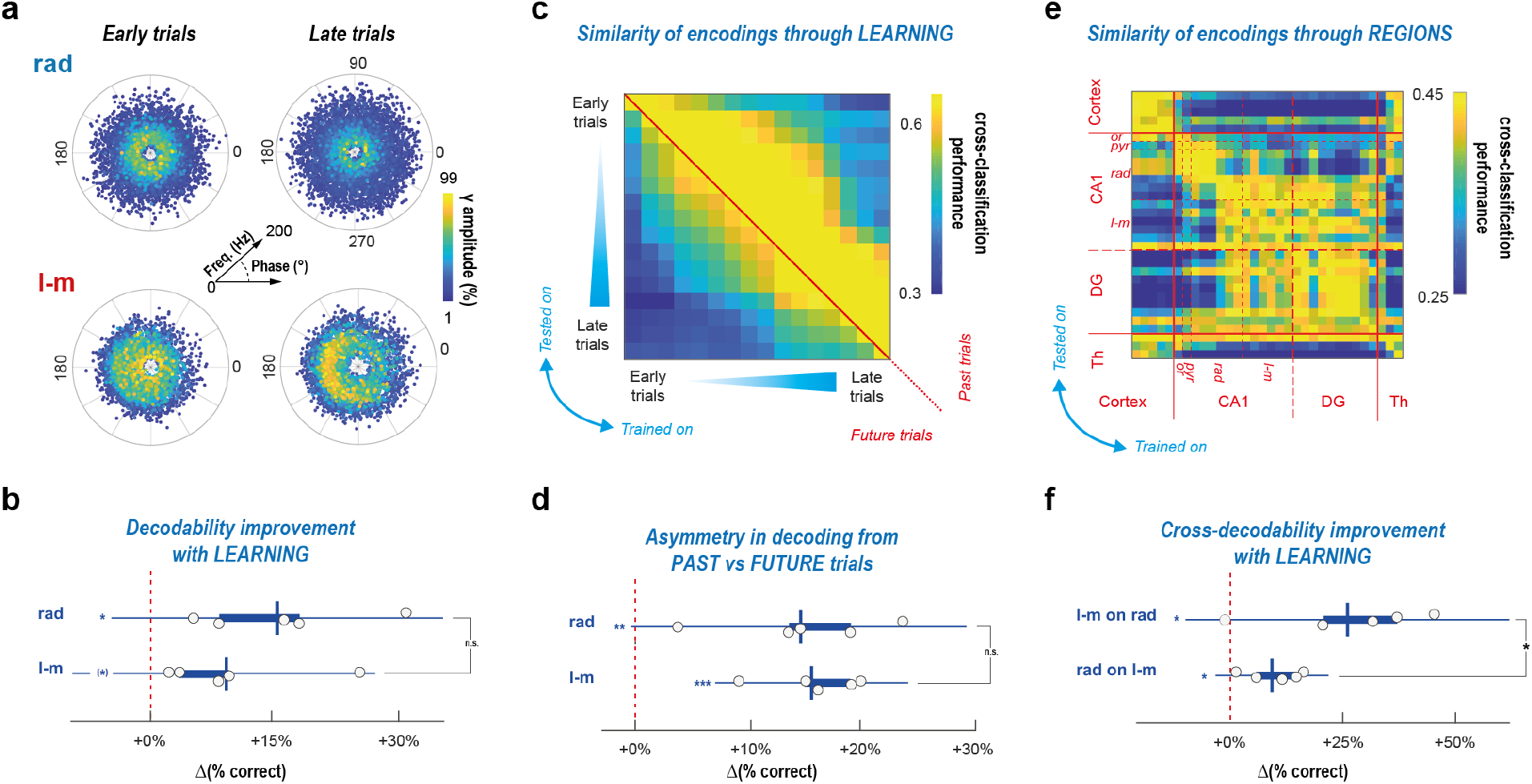
Relations of gamma elements with behavior depend on training and anatomical layer. **a,** Polar scatter plots of individual gamma elements distribution (as in Figure 1d), separate for early (days 1-3) and late (days 8-10) trials. Wide diversity of gamma elements features is observed at all stages of task learning (here for mouse #3; see Extended Data Figures 2, 5a and 11 for more details and all mice). Although remaining complex, the distributions are however evolving with learning. In boxplots: dots denote relative performance improvement between late and early trials, for different mice; boxes, IQRs; horizontal line, sample mean; whiskers, 95% sample c.i.; *, p < 0.05; **, p < 0.01; ***, p < 0.001 after Bonferroni correction; symbols in brackets denote significance only before Bonferroni correction. One-tailed t-tests are used for both comparisons of: samples with chance level; and between samples. **b**, The average performance of decoding maze location (average over all classes, for reference channels in *rad* and *l-m*) is higher for late than for early trials, as revealed by boxplots of percent performance improvement for representative channels in both *rad* and *l-m* layers (p < 0.013 for rad and p < 0.04 for *l-m*). **c-d**, We performed cross-classification analyses, training classifiers to decode maze location from gamma ensembles in a range of trials and using them to extract information from other trials in past or future time ranges. **c,** The resulting cross-prediction error matrix (here for a representative *rad* channel for mouse #3; see Extended Data Figure 12a for *l-m* layer and other mice) is asymmetric with respect to the diagonal, indicating that classifiers trained on future trial ranges can decode information from past trial ranges better than in the opposite direction. **d,** This asymmetry is quantitively confirmed by positive percent difference between performances in past-on-future or future-on-past prediction directions (positivity of the increment, p < 0.006 for *rad* and p < 0.0006 for *l-m*). **e-f,** We also studied cross-classification between different recording locations and its variations along task learning. **e,** The cross-prediction error matrix (all trials, mouse #3; see Extended Data Figure 12b for other mice) displays a block structure, indicating that different anatomical locations have alternative types of gamma elements -to- behavior inter-relations. Hippocampal and non-hippocampal channels form different blocks. Within hippocampus CA1, superior rad and *l-m* channels belong as well to different sub-blocks. **f,** These anatomically-organized patterns of inter-relations evolve along task learning, as revealed by increased *l-m* vs *rad* cross-decodability in late with respect to early stages (percent improvement of cross-prediction performance; p < 0.016 for both *l-m-on-rad* and *rad*-on-*l-m* cross-prediction directions). The improvement in cross-predictability across learning was larger in the *l-m-to-rad* than in the *rad-to-l-m* direction (significance of difference, p < 0.042).

We then compared the complex ways in which gamma element variations reflected maze location by adopting a cross-classification approach. Classifiers trained on trials within a specific training trial range were used to predict maze location on trials from another testing trial range, and the obtained fractions of correct prediction were compiled into cross-prediction performance matrices (see **Fig. 4c** for a representative *rad* layer example and **Extended Data Fig. 12a** for *rad* and *l-m* layers in all mice). The obtained matrices of cross-prediction performance across trial ranges were characteristically asymmetric: the larger upper than lower triangular parts indicate that a classifier trained in one trial range can better predict location from past rather than future trial ranges (cf. more yellow above the diagonal in **Fig. 4c** and **Extended Data Fig. 12a**; see also **Fig. 4d** for a quantification). The performance of decoding yet dropped when the training and testing trial ranges were separated by a timespan too large (cf. blue zone at the upper right corner in **Fig. 4c** and **Extended Data Fig. 12a**). We interpret these findings as an indication that the complex mapping of maze location by gamma ensemble features is not frozen but smoothly evolves through time. However, the drift of this mapping is specifically shaped by previous experience, hence the existence of an “arrow of time” in cross-trial decodability.

## Complex gamma ensembles are spatially organized

We adopted an analogous cross classification approach to compare the mappings of maze location by gamma ensembles recorded at different anatomical locations. **Fig. 4e** shows a representative matrix of the fraction of correct predictions obtained when training a classifier on gamma elements recorded on a channel and testing it from gamma elements recorded on another channel (see **Extended Data Figure 12b** for all mice). This matrix displays a hierarchical block organization. Classifiers trained on channels within the hippocampus can decode maze location from other hippocampal, but not extra-hippocampal, channels (and vice versa). Furthermore, within CA1, at least two blocks can be distinguished including channels located within the *pyr* and upper *rad* layers, and lower *rad* and *l-m* layers, respectively. Cross-decodability between classifiers was high between channels from the same block, but low with extra-block channels, indicating that at least two types of CA1 gamma ensembles exist, differentially modulated by behavior despite their large overlap in frequency and phase distributions.

We then repeated this spatial cross-classification analysis but separately for earlier and later trials along the learning of the task (**Extended Data Figure 12b**). We found that the cross-decodability between the *l-m*-like and upper *rad*-like channel blocks increased in later trials. In general, cross-decodability increased with task learning between all channels. However, this was particularly noticeable for classifiers trained within the *l-m*-like channel block as they gradually improved in decoding the maze location from gamma elements recorded in the upper *rad*-including channel range (**Fig. 4f**). Such results may suggest a convergence of current sensory representations conveyed by entorhinal inputs to the *l-m* layer, onto internal model representations, provided by CA3 inputs to the *rad* layer (see *Discussion*).

## Discussion

Using machine learning-based decoding of electrophysiological recordings during a behavioral task, we showed that *in vivo* hippocampal gamma oscillations are not well described by sharply distinct narrow-band modes. On the contrary, at every CA1 channel, we observed broad distributions of gamma frequency and theta-phase of appearance, largely overlapping between distinct anatomical layers (**Fig. 1**, **Extended Data Figs. 2-6**). Hippocampal oscillations would thus be better described in terms of a collection of complex “gamma ensembles”, i.e. collections of transient oscillatory events that, despite their heterogeneity in phase and frequency and their generally low amplitude, are distinctively modulated by both behavior and learning. This diversity of gamma oscillations is widespread and common to all analyzed datasets (**Extended Data Figs. 3**). However, even studies reporting this diversity and calling for abandoning a strict dual gamma band view chose implicitly to emphasize a minority of high amplitude oscillatory events^2,13,14,22^, more localized in phase and frequency, which are the ones dominating average pattern analyses^3^ but are not well representative of the transient oscillatory dynamics most frequently observed when not averaging.

Here we argue that the haphazard heterogeneity of frequencies and phases presented by low amplitude gamma oscillatory elements can have natural mechanistic explanations. Our computational model shows that a balanced excitatory-inhibitory recurrent circuit can generate oscillatory events with fluctuating and diverse frequencies –and thus a complex gamma ensemble– if parameters are liberally chosen to be in a broad fringe in proximity but still below a transition to partially synchronized collective oscillations ^23^. This contrasts with many previous models ^24,25^ which operated in more synchronous regimes: their narrowly tuned rhythms, although matching prescribed target frequencies based on average oscillatory patterns observed *in vivo*, cannot indeed render the strong variability observed for individual gamma elements. Nevertheless, even these exceedingly synchronous models already showed that the frequencies of gamma oscillations are not hardwired but result from a complex interplay of network-level factors and could thus be dynamic. Dynamics at the fringe-of-synchrony, like the one produced by our model, are intrinsically collective, robustly emerging from local network interactions without the need of neuronal properties fine-tuning. Despite their apparent stochasticity, oscillatory transients at the fringe-of-synchrony can still route information and self-organize into well-defined phase-relations with other coupled populations, at least during short-lived events with high power^4^. Such predicted phenomenology is well compliant with the observed properties of inputs received at the *l-m* and originating mostly from the entorhinal cortex, which are a mixture of low-power gamma elements scattered in phase with higher-power elements concentrated around a specific phase (**Fig. 2**, **Extended Data Fig. 2 and 6**). On the contrary, inputs received at the *rad* layer are scattered in phase at all power levels, suggesting that the generating populations in CA3 may be tuned in an even more asynchronous regime, only transiently “ringing” as an effect of filtering noise^26,27^.

We also additionally proved that fluctuating gamma elements convey information about behavior. Maze location can be decoded with comparable accuracies from both low and high gamma amplitude events, showing that behavior is not only modulating the extreme gamma elements which dominate ordinary averages but also those within the distribution bulk and ignored by more conventional analyses. A possible explanation for the success of our decoding is that oscillatory features such as phase, amplitude and frequency or cross-frequency relations provide a code for location, or, in other words, that their modulations have a representational meaning. This hypothesis is implicit in some previous studies that already performed decoding of location based on hippocampal signals^28^. Another possibility is that oscillatory fluctuations are just indirect signatures of other codes, relying on cell ensemble firing^29^ or the dynamic selection of internal attractors or assemblies^30,31^. Changing neuronal correlation and firing may translate into broadband deformations of the extracellular field power spectrum shape^32^, thus explaining why information can be decoded even from oscillatory events at frequencies remote from spectral peaks.

Previous studies showed that increasing speed was associated with modulations of gamma amplitude, frequency or theta-phase, often with elaborate nonlinear relations, possibly dependent on learning^32^. The difficulty to identify relations between individual oscillatory features and speed may reflect the essentially synergistic, and thus conditional, nature of the mapping between gamma element variability and exploration behavior (**Fig. 3e-g**). Some aspects of the performance of our decoders may well hint at an influence of speed on the classifier decisions. For instance, it was easier to correctly identify presence in the target arm when speed was high, and in the reward field when speed was low. Similarly, gamma elements during fast movement in other arms tended to be misclassified as occurring in the target arm (cf. **Extended Data Fig. 9b**). Nevertheless, decoding performance remained well above chance level for all speed quartiles. Furthermore, the redundancies of individual gamma elements feature with speed (**Fig. 3h**) and the unique information about maze location conveyed by speed (**Fig. 3g**) both remained very low. Maze location affects the features of gamma elements in complex and synergistic ways, particularly in the case of the *l-m* gamma ensemble, whose synergistic mutual information levels where systematically larger than for the *rad* (cf. **Extended data Figure 10**), possibly reflecting richer and higher-dimensional encoding schemes by the activity of entorhinal cortex than of CA3 circuits. It is likely that superior decoding, potentially allowing a finer identification of location beyond our rough subdivisions, becomes accessible when going beyond individual gamma elements to consider combinations of them (either temporal sequences or co-occurrences across layers).

Despite a substantial overlap in the frequency distributions of gamma elements at all layers, our cross-decodability analyses could yet reveal the existence of at least two types of gamma ensembles, corresponding to distinct blocks in the matrices of **Fig. 4e** and **Extended Data Fig. 12b**. Gamma elements recorded within the *l-m* (and the dentate gyrus) were modulated by maze location in very similar ways, but differently from channels within the more superficial *rad*. Remarkably, it is at the *algorithmic* level^33^ of how behavior-related information is encoded that we can recover a clear distinction between the *rad* and *l-m* layers that was not so evident at the level of frequency and phase distributions. Our results provide a further confirmation that spatial location and navigation behavior are differently represented by the entorhinal cortex and CA3 circuits, serving as the input sources for functionally distinct gamma ensembles. These representations are not fixed but evolve through the learning of the task, and more information about location can be decoded from late trials (**Fig. 4b**). Such improvement is paralleled by (and may be attributed to) an increase in the cross-decodability between layers, meaning that the nature of the representation is dynamically transformed. Decoders able to read sensory-related representations at layers receiving an entorhinal input become increasingly able to equally read model-based representations at layers innervated by CA3 (**Fig. 4f**). In other words, the grammar of sensory-related inputs, parsed by our decoders, becomes more and more compliant with the one of internal models. This result finds a natural interpretation within a predictive brain framework, since, with the learning of an internal model, the activity of sensory-processing regions shifts toward representing model-based inferences, beyond a passive encoding of external evidence^34^.

Finally, our results show that there is not a sharp distinction between naïve and expert types of location representations, but rather that these representations are smoothly adjusted through time as an effect of idiosyncratic experience. Our analyses indeed reveal an “arrow of time”, with present decoders able to read out information from past gamma elements but not yet future ones (**Fig. 4c-d**). Each mouse has a different history of learning and, thus, potentially, a different way of coding rich individual behavior into differently organized but invariantly complex languages based on gamma ensembles (eventually, cross-classification between mice was not significant).

To conclude, hippocampal theta-gamma activity is more diversified than a limited number of narrow frequency bands used by afferent generators at specific phases of the ongoing theta oscillation. At first sight, this variety may seem to threaten prominent views in which information from the two main afferents is conditionally routed and disentangled at the neuronal level thanks to their distinct preferential frequency and theta-phase^14,22,35,36^. In these views, indeed it is a precise temporal and spectral separation of inputs which allows different structural pathways to mediate distinct cognitive functions^11,37^. Here, we show that such precise separation in frequency and phase most of the times does not occur. Yet, the diversity of gamma ensembles is not mere noise as random-like gamma elements, usually discarded, still allow the successful decoding of behavior. Furthermore, their complex conditional distributions, harnessed by our machine-learning classifiers, are still meaningfully coupled to both anatomy and learning. By emphasizing the relevance of low power events with “misbehaving” phase and frequency, our results suggest that system’s function may rely on the self-organized coordination between noisy and weak oscillatory bursts^4^ rather than on rigid architectures with precisely tuned oscillations.

## Methods

### Animals and surgery

#### Subjects

Five adults male CD1 mice (~3-month-old at the time of surgery) were housed in individual cages post-surgery, under a 12-h light/dark cycle (light at 8:00 A.M.). They had water and food *ad libitum* till the start of the habituation period; they were then water-restricted (2h-daily access, ~2h after testing) for the entire duration of the experimental protocol. All experimental protocols agreed with the European Committee Council directive (2016/63/UE) regarding animal experimentation and were approved by the French Ministry of Research (APAFIS#20388-2019042517013497).

#### Surgery

Animals were anesthetized with isoflurane during the entire surgery. Linear silicon probes with either 16 or 32 channels (50 μm-spacing; A1x16-3mm-50-177-CM16LP or A1x32-6mm-50-177-CM32, Neuronexus, Ann Arbor, USA) were chronically implanted through the CA1-DG axis of the right dorsal hippocampus (AP: 2.06 and ML: 1.3 from bregma; DV: 1.7 from the dura). They were covered with DiI stain (Invitrogen Molecular probes, USA) before insertion. Two screws were positioned in posterior and anterior portions of the skull, serving as ground and reference electrodes, respectively.

#### Histological procedures

The mice were perfused with 0.1 M PBS followed by 4% paraformaldehyde in PBS solution with added heparin (25 kUI). Brains were postfixed for 24 h in 4% paraformaldehyde before being cryoprotected in 20% sucrose solution for 48 h. They were then frozen in isopentane and sliced into 40-μm coronal sections. Implantation sites were visualized through a fluorescence microscope (Zeiss) thanks to the Dil stain.

### Behavioral apparatus and protocols

#### Eight-arm radial maze

The radial arm maze consisted in a central platform (52-cm diameter) from which eight identical arms (55 x 10 cm) expanded, separated by a 45-degree angle. Each arm was surrounded by a 3-cm high wall. A shallow circular recess at the end of each arm could hold the reward (75 μL of 5%-sucrose solution). The maze was situated 65 cm above the floor in a room displaying numerous distal visual cues that remained in position for the entire duration of the experiment. Mice were transferred from their home cage to the maze using an opaque box (start box: 20×10×15 cm).

#### Habituation to the apparatus

One week after surgery, mice were habituated to the experimental apparatus and the experimenter. A recording cable was plugged on a permanent basis to the headmounted pre-amplifier so the mice could get used to its presence and weight. The animals were then handled by the experimenter for a few days before starting the habituation *per se*. This period consisted in transferring the mouse from its home cage to a single arm removed from the maze and placed elsewhere in the room. The mouse had to wait for 20 s in the start box (positioned at the entry of the arm) before the door opening. The aim was to reach the other end of the arm to consume the reward. This was repeated for three to five days, with five to eight trials a day (or until the mouse was not showing clear signs of anxiety). Mice were then exposed to the radial arm maze for two days during a daily 10-min trial in which the start box was positioned at the center of the maze and opened after 20 s. Every arm was reinforced only once per trial to promote exploration of all arms across both days. The inter-trial interval was of five min, during which the apparatus was cleaned with 35% ethanol.

#### Arm-to-Arm task

In the Arm-to-Arm (ATA) task, mice must find the rewarded arm, the same across the 10 days of training (~24 h between sessions). The four daily trials start each from one of the four possible departure arms (two and three arms away from the target arm, both left and right; identical across sessions to ensure a constant distance to target) following a pseudo-random order to promote allocentric navigation. Hence, the animals wait in the start box positioned at the end of one arm for 20 s before opening of the door. It then has up to three min to find the rewarded arm otherwise the trial is stopped. The inter-trial interval was of five min, during which the apparatus was cleaned with 35% ethanol. On the 11th day, a five min probe test is carried out to assess the animal’s spatial reference memory: the mouse is released from a new departure arm (opposite to the target) and no reward is available. The mouse is considered to have learned the reward location if it either visited more often or spent more time in the target and its two adjacent arms than chance (proportion: 0.125).

### Electrophysiological recordings and analysis

#### Recording and preprocessing

The electrophysiological activity was recorded with an Intan recording controller (RHD Recording Controller, Intan Technologies, USA). The signals were amplified 200x, recorded whole-band (0.1-10 kHz) and digitized at 20 kHz. They were synchronized with a video system tracking the position of the animal at 20 Hz (Imetronic, France). The basic pre-processing of the LFPs included the removal of both slow variations and 50-Hz (and harmonics up to 200 Hz) electrical noise (Chronux Matlab toolbox^38^), artefact correction^39^ and finally downsampling to 1 kHz.

#### Anatomical localization of the electrodes

Each electrode was assigned to an anatomical hippocampal layer depending on its distance from the hippocampal fissure along the estimated probe position in the histological slice. The theta power from each electrode was calculated by a group of complex Morlet wavelets (1-14 Hz by 1-Hz steps; 2-s duration; number of cycles linearly dependent on frequency, between 2 and 4 cycles) on the LFPs filtered for theta range (4-12 Hz; zero-phase digital filtering using a finite impulse response filter of order = 256). The fissure was located at the peak of the Gaussian fit of the theta power curve, possibly between two electrodes.

#### Signal decomposition

For further analyses, instead of using a classic passband filter, we used an unsupervised, nonlinear and non-stationary technique to isolate the dominant oscillations present in the LFPs in time, amplitude and frequency: the Empirical Ensemble Mode Decomposition (EEMD ^40^). The resultant components, termed Intrinsic Mode Functions (IMFs), can then be summed to recompose the original signal. Hence, to filter the LFPs in either theta (4-12 Hz) or gamma (30-250 Hz) frequency range, we summed the IMFs whose mean of the Hilbert-derived instantaneous frequency fell within the relevant range, thus obtaining a theta and a gamma composite LFP signals. For every trial, LFPs were decomposed independently for the period of actual navigation, that is, from when the animal is about to navigate in the maze (hence excluding start box or behavioral inactivity periods sometimes following the box opening) to up to 5 s following arrival to reward (or trial end if the animal did not find the water). Ten IMFs were requested, resulting from the average of 2000 iterations with added noise (input noise level of 0.3 except for some trials from two mice [mouse #3 and #4] needing 0.8 to satisfactorily alleviate mode mixing). To reduce confounds from potential theta harmonics, we started our gamma range at 30 Hz^2^ unlike some previous reports of a lower bound at 25 Hz (see^41^ for a recent review). Note that, to contrast our results with established methods (**Extended Data fig 3**), we also processed the signal using a finite impulse response filter combined with a zero-phase filtering for both theta and gamma bands or with an independent component analysis (KD-ICA algorithm within ‘ICAofLFPs’ Matlab toolbox^42^) instead of the EEMD decomposition.

#### Theta cycles identification and selection

Theta cycles were identified on the theta LFP composite from the closest channel to the hippocampal fissure. Using the fissure as a theta reference offers larger, more defined theta cycles but implies an inverted theta phase compared to theta recorded within the CA1 pyramidal and oriens layers. Peaks in the signal were identified as the start/end of each candidate cycle. The trough was determined as the point with the lowest amplitude between two consecutive peaks, and the flanks, as the points at half-amplitude between the trough and these surrounding peaks. The theta phases (0-360°; peak = 0/360°) were obtained by linear interpolation within each quadrant formed by the starting peak-descending flank (0-90°), the descending flank-trough (90-180°), the trough-ascending flank (180-270°) and the ascending flank-next peak (270-360°). This waveform-derived phase determination is more respectful of the theta waves asymmetry than the one from the Hilbert transform^43,44^ although we compared both methods (**Extended Data fig 3**). Note that EEMD-based composite signals are supposed to better respect the wave asymmetry than classic filters^13^. To be selected for analysis, the candidate theta cycles had to meet the following criteria^13^: a duration compatible with the theta frequency band (i.e., 83 to 250 ms) and a sufficient power (amplitude of the envelope of the theta LFP composite signal at the cycle start, mid and end points superior to the envelope of the 1-4 Hz infra-theta LFP composite signal). They further needed a coincident video sample to determine the animal position in the maze at that time.

#### Amplitude of theta cycle-nested gamma

To lessen volume-conducted activity, the amplitude of gamma oscillations was calculated on the current source-density (CSD) signal derived from gamma LFP composites as previously described for LFP^36^. CSD at a given time point *t* was calculated as follows:

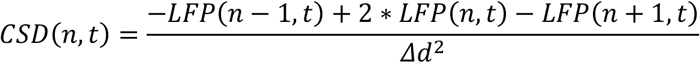

where LFP_(n,t)_ is the gamma LFP composite recorded at the electrode *n*, LFP_(n-1,t)_ and LFP_(n-1,t)_ are the gamma LFP composites from electrodes directly above and below, respectively, and *Δd* is the distance (in mm) between contacts.

The continuous amplitude of the CSD signal, used as an instantaneous metric of power, was then obtained for each channel using complex Morlet wavelets convolution (0.5-s duration; from 15 to 200 Hz by 5-Hz steps and assessed by a number of cycles linearly dependent on the wavelet main frequency, between 6 and 20 cycles). The portion of this convolution corresponding to the time of each theta cycle was then isolated^13,14^ and the CSD amplitude for each gamma frequency was averaged per theta phase (10° phase bins). Hence, the gamma spectral contents of each theta cycle were summarized in a ‘snippet’ (38 x 36 matrix: frequency x theta phase bin).

#### Gamma bouts detection

Within each individual theta-cycle, we extracted gamma elements as patches of locally higher gamma composite power in the CSD spectrogram. To identify these patches, we treated single-theta cycle spectrograms as color-scale images and binarized them, assigning to pixels with a gamma composite power larger or lower than a fixed threshold black or white color, respectively. We then used a standard flood-fill algorithm^45^ to identify connected components within the binarized spectrogram image, each corresponding to a potential gamma patch. Since the number of connected components depend on the applied threshold, we decreased systematically the threshold starting from a value equal to the maximum power value within the original spectrogram. When reducing the threshold, more image pixels rise above threshold and the number of connected components tend to increase, apart from a few exceptions (see below). The scanning of decreasing threshold values stopped when a maximum (arbitrarily chosen) number of four connected patches (and thus gamma elements per theta cycle) was identified. Cases could arise in which the addition of new black pixels to the binarized image caused patches disconnected at higher threshold values to finally merge. However, such patch fusion should be prevented, as the merged patch includes multiple and distinct power peaks. We thus added tracked record of the components’ extensions immediately prior to merging, storing them as separated. Another special case needing ad hoc handling was the one of components located at the boundaries of the theta cycle and therefore potentially extending across two contiguous theta-cycle. To avoid double counting of a same component (detectable in both the cycles across which it is split), we thus parsed simultaneously neighboring theta cycles, to identify the complete extension of cross-cycle boundaries patches and count them only once (assigning them to the cycle over which the strongest amount of power was located).

After determining connected components segregated from the background and correcting for patch fusion and double counting, we then computed for each retained component the following gamma element features: gamma amplitude, frequency and theta-phase of occurrence. Each pixel within a component was associated to a specific power, frequency and phase triplet of values (respectively, the color, the vertical and the horizontal coordinate within the single-theta cycle spectrogram image). The gamma element power was evaluated as the average power over all pixels within a connected component. The gamma element frequency and phase were then determined as the average among the frequencies and phases of the pixels within the component, weighted pixel-by-pixel by the pixel power.

We also compute the associated theta wave instantaneous frequency (inferred from cycle duration), amplitude (average voltage difference between the trough and the two adjacent peaks) and asymmetry (rise – decay ratio).

All gamma elements were appended to a list for each trial and channel, which was then filtered to exclude the top and lowest 1 % amplitude gamma elements. Analogously, we excluded some gamma elements occurring in theta cycles coincident to unlikely large running speed (> 100 cm/s).

### Dimensionally reduced representations of gamma elements

We used a standard t-Stochastic Neighborood Embedding (t-SNE) algorithm^16^ to create bidimensional representations of the diversity of gamma elements. This algorithm guarantees that distance inter-relations between data-points in the source high-dimensional space are preserved as much as possible in the target bidimensional space representation. The projection was learned for all gamma elements simultaneously (all layers and trials), then different group of elements could be shown in different panels (see. **Fig. 1f-g** and **Extended Data Fig. 5**) filtering a same common and frozen projection. We used standard hyperparameters (perplexity = 30, no exaggeration) with an approximated Barnes-Hut algorithm. We used a Euclidean distance metric except for the theta phase of gamma appearance where circular distance was used.

### Computational model of gamma elements generation

#### Model definition and parameters

We considered a network composed of N=2000 quadratic integrate and fire (QIF) neurons^20^, 80% of them excitatory (E) and 20% inhibitory (I). The membrane potential 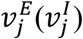 of each excitatory (inhibitory) neuron j obeyed the following differential equations:

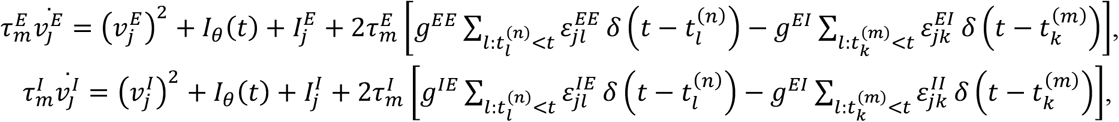

where 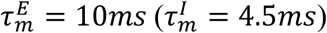 is the excitatory (inhibitory) membrane time constant and 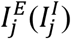 the neuronal excitability encompassing single neuron characteristics as well as synaptic drives originating from other neural regions and acting on the excitatory (inhibitory) neuron *j*. The input term *I_θ_*(*t*) is a forcing current periodically modulated at a θ-like frequency of 10Hz and *g^αβ^* the synaptic coupling strength between a post-synaptic neuron *s* in population *β* and pre-synaptic neurons in population α, with [α,β] being either E (excitatory) or I (inhibitory). The connectivity matrix elements 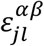 are equal to 1 (0) if a connection from a pre-synaptic neuron *l* of population β towards a post-synaptic neuron *j* of population α, exists (or not). Furthermore, 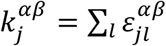 is the number of pre-synaptic neurons in population β connected to a neuron *j* in population α, or, in other terms, its in-degree restricted to population β. The emission of the *n*-th spike emitted by neuron *l* of population α occurs at time 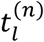 whenever the membrane potential *v^α^* crosses threshold for firing, while the reset mechanism is modeled by resetting *v^α^* to a rest value, immediately after the spike emission (see ^46^ for details on threshold and reset in QIF neuron model). For the sake of simplicity, we assumed synapses to be fast and synaptic transmission instantaneous, therefore the post-synaptic potentials were modeled as δ-pulses without any delayed activity. Connectivity within the E and I populations was random and quenched, with in-degrees *k^αα^* distributed according to a Gaussian distribution with mean *K^αα^* and with a standard deviation *Δ^αα^*, this latter parameter measuring the level of structural heterogeneity in each population. We chose here to set *K^EE^* = *K^II^* ≡ *K*, providing a common scale for the strength of local connectivity in the model. As a further simplification (suitable for potential mean-field reduction not explored in this study), we then assumed that the E and I populations are globally cross-coupled, i.e. 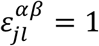 if *α* ≠ *β*. The neuronal excitabilities 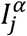 were distributed according to a Gaussian distribution with mean 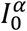 and standard deviation *D^α^*. The DC currents and the synaptic coupling were rescaled with the median in-degree as 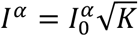 and 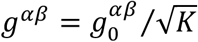 to obtain a self-sustained balanced dynamics for *K* → ∞ ^23,47,48^. The structural heterogeneity parameters were rescaled as 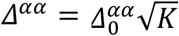 in analogy to Erdos-Renyi networks ^23^. We employed, unless stated otherwise, the following values of the parameters: 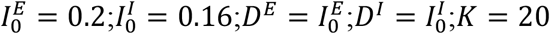; 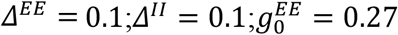; 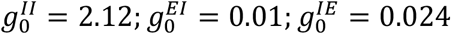. The θ-forcing was assumed to be perfectly sinusoidal, as 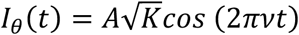, with *v* = 10*Hz* and *A* = 0.042.

#### Simulated Local Field Potential

The Local Field Potential was modelled as LFP= (|*I_A_*|+|*I_G_*|), which is the sum of the absolute values of AMPA and GABA currents impinging on pyramidal cells, following^49^. The global currents *I_A_* and *I_G_* were the linear sum of contributions induced by single pre-synaptic spikes, each represented as a combination of two exponentially decaying functions. This representation can be obtained using auxiliary variables *x_Aj_,x_Gj_*. The time evolution of AMPA and GABA-type currents of neuron j were thus described by the following ordinary differential equations:

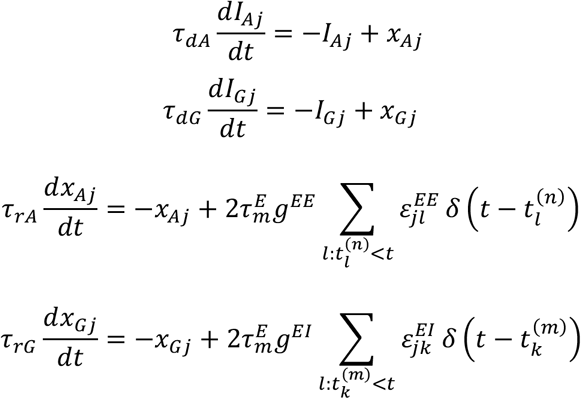

where *τ_dA_*(*τ_dG_*) and *τ_rA_*(*τ_rG_*) are respectively the decay and rise time of the AMPA-type (GABA-type) synaptic currents. Always following^49^, we high-pass filtered the obtained model LFP signal at 1 Hz with a 4th order Butterworth filter and employed *τ_rA_* = 0.4*ms*; *τ_rG_* = 0.25*ms*; *τ_dA_* = 2*ms*; *τ_dG_* = 5*ms*.

#### Numerical simulations

Numerical simulations of the model were performed with a standard Euler integration scheme with time step δt=0.001ms. Since all disorder in connectivity and conductance is quenched, a deterministic integration scheme can be used as in^46^. Simulations were performed scanning a range of *K* and 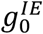 values to explore different dynamical regimes (cf. **Fig. 2 and Extended Data Fig. 7**).

#### Indicators of dynamic regime

After the generation of synthetic LFPs, gamma elements could be extracted from them following the same procedures as for real LFP and CSD signals. We also computed an indicator of power distribution across frequencies, computing spectral entropy. To do so, the power spectrum *P(f)* of simulated time-series was computed and over a range between *f_min_* = 25 Hz and *f_max_* = 125 Hz sampled at *df*=0.1Hz and normalized to provide a density functional. We then evaluated spectral Entropy as the quantity 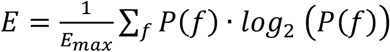, where *E_max_* = *log*_2_(*M*) and *M* = (*f_max_* – *f_min_*)/*df*. Higher values of *E* correspond to higher dispersion of power across gamma frequencies.

We also estimated the ratio between the amounts of high- and low gamma power, estimating the total power in the low (25-50 Hz) gamma band P_low_, and the total power in the high (50-100 Hz) gamma band P_high_. The ratio rgamma (P_low_-P_high_)/(P_high_+ P_low_) indicated whether the LFP is dominated by high gamma (positive r_gamma_), by low gamma (negative r_gamma_), or by an equilibrated mix of the two (r_gamma_ close to zero).

Finally, we computed standard deviation of the mean membrane potentials across inhibitory neurons as a measure of the amplitude of generated gamma oscillations.

The values of spectral entropy, high/low gamma power ratio and standard deviation of potential were computed for simulations of models with different connectivity to study the dependency on them of the obtained dynamical regimes.

### Classifier training

#### General classification approach

We used a supervised classification approach to predict rough location within the maze based on an input vector parameterizing individual gamma elements. The input was given by a six-dimensional vector including general information about the theta cycle (theta-composite amplitude, frequency and asymmetry) and specific features of the considered element (gamma element power, frequency and theta phase of occurrence), computed as described in previous sections. The output was a categorical label, referring to a subdivision of the maze in four sections: *“RewardRF”* (end-field of target arm where reward was delivered); *“target arm”* (including the arm leading to the reward location and the outer area of approach to this same arm, “); *“other RFs”* (arm end field other than the reward field); and, finally, *“other locations”* (including all areas not including in the previous subdivisions, i.e. maze center and generic maze arms not leading to reward).

As multi-class classifiers, we used boosted ensembles of classification trees, limiting the maximum number of decision splits in a tree to 500, and the number of learners in an ensemble to 500 trees. Tree ensembles were fitted using the RUSBoost algorithm^50^, with a slow learning rate of 0.01, to alleviate the problem of output class unbalance (as some classes, as *“target arm”* or *“Reward RF”* are under-represented relatively to others). In this algorithm, random undersampling is applied to training sets to guarantee that each class is represented by close numbers of samples, providing simultaneously capability to learn rare classes and protection against biases due to variations across different conditions (e.g. early vs late trials) of the fractions of samples per class.

#### Classification performance

Classification performance was evaluated both in terms of resubstitution error (error on same data samples used for training) and generalization error (error on data samples not used for training), estimated via 4-fold cross-validation. In generating the random partitions into training and testing sets of elements, beyond output class balancing, we took care to use for testing gamma elements measured in theta cycles not included in the training set, thus conferring protection against overfitting, as covariations in features may subsist among gamma elements occurring within the same theta cycle. The list of theta cycles (and gamma elements therein) available for selection of training and testing pools corresponded to the total list of elements retained for a channel in a mouse, over all the trials (unless otherwise specified, see next section on cross-classification). Different classifiers were trained independently for each different channel. **Fig. 3d-f** and **Extended Data Fig. 9** report average performances over all CA1 channels, as classification performance was shown to have only weak dependency from the layer (cf. Fig. 3c and **Extended Data Fig. 8**). Generally, unless otherwise specified, classification performances (and misclassification rates) are expressed in terms of correct (incorrect) classification fraction, evaluated over all available gamma elements (cross-validation ensuring that prediction on an element was performed in terms of classifiers not having seen this element during training). Despite cross-validated training had access to the whole list of gamma elements extracted from a channel, after training, we could also evaluate classification performance on subsets of gamma elements to assess whether the probability of correct classification depended on various features of the gamma elements fed as input. We thus separated gamma elements according to them belonging to different quartiles of the distribution of different features (from Q4, with top values, to Q1, with the lowest values): the six features of the gamma element descriptive vector (gamma power, frequency and phase, theta amplitude, frequency and asymmetry), as well as motion speed (averaged over the time range of the considered theta cycle); and computed fractions of correct classification separately over each distribution quartile.

#### Classification with alternative input features

We also trained classifiers using alternative reduced sets of input features. Instead of using the full six-dimensional descriptive vectors of gamma elements as input vector (as in the *“theta + gamma”* classifiers just described), the *“theta-only”* (or *“gamma-only”*) classifiers were trained just in terms of the three theta (or gamma) features entries.

#### Classification of alternative arm and in probe trials

We also used an alternative set of output labels in which the *“Target arm”* section of the maze was merged with the “other locations” section, but in which an *“Alternative arm”* was considered instead as a separate section, with the same extension of the Target arm zone (arm plus outer arm approach zone) but including an arm different from the one leading to reward. This alternative arm was chosen to be opposite to the target one. In this alternative zone labelling, the *“reward RF”* zone was left unchanged, i.e. it still included the reward location (and was therefore not contiguous to the *“Alternative arm”* zone). Classifiers were trained on this new labelling in a completely independent way from the classifiers trained on the original labels.

Probe trials were not used for training, but locations prediction was performed using classifiers trained over control trials, with the ordinary output zone labeling (i.e., including the *“Target arm”* and not the *“alternative arm”* zone).

### Cross-classification

#### General approach

Once trained, a classifier serves as an implicit model of the distribution of gamma ensembles in relation to behavior. Changes of the ensembles-to-behavior relation across conditions or channels can be studied using a cross-classification approach, in which classifiers trained on a sample are evaluated on a different sample. Preserved or decreased performance levels will then denote, respectively, similarity or dissimilarity or relation to behavior.

#### Cross-classification through learning

To study evolution of the ensemble-to-behavior mapping across task learning by the mice, we selected gamma ensembles over subsets of trials only. Specifically, we sorted all trials available from the earliest to the latest and compiled a table of how many gamma elements each trial provided on average over all channels. We then defined two *“early”* and *“late” trials* ranges, including trials with ordinal number respectively smaller and larger or equal than a pivot trial number. This pivot trial number was chosen such that the cumulative sums of gamma ensemble counts per trial over the early and late ranges were as close as possible between them. The early and late trial range specifications were therefore adapted to the actual behavioral history of each mouse. Furthermore, the early and late trial ranges usually included unequal numbers of trials, as maze exploration is faster in later than in earlier trials and, consequently, individual late trials usually contribute smaller counts of gamma ensembles.

We then adopted a finer subdivision of trials when constructing the cross-classification matrices of **Fig. 4c** and **Extended Data Fig. 12a**. Once again, we ordered trials and grouped them into smaller window ranges such that the cumulative sum of gamma ensemble counts for the trials included in each window was as close as possible to 3000 elements. Every window included all trials with ordinal number between the ones of a start and stop trials. Windows could have an overlap, but two consecutive windows could not have the same start and stop trials. Different windows generally included different numbers of trials, with windows at earlier times being generally narrower than windows at later times.

Classifiers were then trained over just the early or late range of trials, or, yet, just trials within a specific learning window, using the same class-balanced, cross-validated approach described in previous section. The partial datasets were randomly downsampled to exactly include the same number of elements (as the numbers of elements provided by early and late trials ranges or by different windows were close between them, but not identical). Although cross-validation was still used in training, it could not be systematically used in evaluating cross-classification performances, as the original and checking datasets did not include the same theta cycles and partitions generated for the one was thus invalid for the other. Therefore, in the cross-classification performance matrices of **Fig. 4c** and **Extended Data Fig. 12a**, we reported average resubstitution error along the diagonal and, in off-diagonal entries, direct average performance on the considered checking dataset.

The improvement of decodability in late relatively to early trials (**Fig. 4b**) was evaluated, for each mouse, as relative percent difference between cross-validated performance, averaged over all classes, of classifiers trained just on late or early trials (for a representative channel in the middle of rad or of the *l-m* layers). The performance asymmetry in classifying past vs future trials (**Fig. 4d**) was evaluated as relative percent difference between the averages of the upper and lower triangular parts of the cross-classification matrices of **Fig. 4c** and **Extended Data Fig. 12a**.

#### Cross-classification through channels

Classifiers trained on a channel were used to extract location based on gamma elements of another channel. We computed cross-classification performance across channels based on the whole set of available trials (**Fig. 4e** and **Extended Data Fig. 12b**) and also based on just trials in the early and late ranges. For a better comparison with cross-classification analyses across trials we once again computed cross-classification performances and relative percent variations in terms of resubstitution and direct checking errors. Cross-classifiability between rad and *l-m* layers was evaluated averaging cross-classification matrix entries in blocks delimited by channel ranges matching the different layers. Note that some uncertainty exists at layer edges, as electrodes could slightly move from one day to the next, causing some of them to transit above or below the depth delimiting two layers. A channel was thus included in the block average only if it belonged to a specific layer in at least three quarters of the trials used to build the classifier.

### Information-theoretical analyses of element features to behavior relation

#### Mutual Information between pairs of features and maze location

To study the nature of the relation existing between different descriptive features of the gamma element and maze location, we complemented decoding by machine-learning classifiers with information theory analyses ^51^ and computed mutual information between pairs of input variables and simultaneously visited maze location. We used a rough estimation of the probability distributions of input variables, quantizing them into four unequal bins, matching the distribution quartile limits. By replacing feature values by their quartile label in the feature distribution, we then automatically maximized single variable entropies, as entropy for discretized variables is maximal for uniform distributions. Output labels were already categorical and in a number of four, corresponding to the four rough maze sections previously described (Reward RF, Target Arm, Other RF, Other locations). For each pair of quantized input features *f* and *g* and output maze location labels *L*, we computed over the list of all gamma elements for representative channels in rad and *l-m* layers the joint normalized frequency histogram *P(f,g,L)* and, out of it, the total mutual information that the pair of inputs (*f,g*) carries about the output *L*:

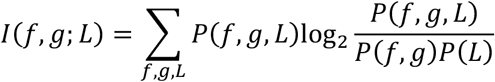

normalized by the total entropy *H*(*L*) = – ∑_*L*_ *P*(*L*)log_2_*P*(*L*) of the output variable (to quantify the fraction of location information carried by the pair of input features).

#### Partial Information Decomposition

We then decomposed this total mutual information using the Partial Information Decomposition (PID) framework^21^ into: *unique* fractions of information, i.e. information that *f* (or *g*) carry about *L* but that *g* (or *f*) don’t carry; a *redundant* fraction of information, i.e. information that both *f* and *g* carry about *L*; and a *synergistic* fraction of information, i.e. information that neither *f* or *g* alone carry about *L* but that they carry when jointly considered. We evaluate the synergistic information of *f* and *g* relative to *L* as:

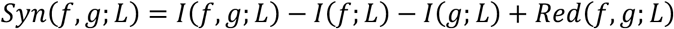

where *Red*(*f, g*; *L*) is the redundant fraction of information and must be added back once because twice removed from the total mutual information *I*(*f,g*; *L*) when subtracting *I*(*f*; *L*) and *I*(*g*; *L*), mutual information of *L* which just *f* or just *g*. To estimate redundancy, we use the so-called Minimal Mutual Information ansatz, under which the redundant information fraction is made to correspond exactly to the minimum between the two individual mutual information terms, i.e. *Red*(*f,g*;*L*) = min [*I*(*f*;*L*),*I*(*g*;*L*)]. In this way, the unique information carried by the least informative of the two variables (say, *g*) is set to zero, and the remaining difference equated to the unique information carried by the most informative variable, i.e. *Unique*(*f*; *L*) = *I*(*f*;*L*) – *I*(*g*; *L*). Unique, redundant and synergistic fractions of the total mutual information can also be normalized by the entropy of the stimulus. **Fig. 3g** shows average total mutual information with location and decomposed fractions, averaged over all pairs of input features including a specific reference feature. Besides the six features describing gamma elements we also considered pairs of inputs including motion speed *V* as input variable, discretized in a quantile-based way as the other features. We also analyzed separately the four gamma elements extracted out of each theta cycle, ranking them in decreasing order of power, to reveal whether the informative content of elements concentrated on the strongest power elements or was uniform across stronger or weaker gamma power elements. Details about the decomposition for specific pairs of features are shown in **Extended Data Fig. 10**.

#### Redundancy with speed

The general dependency of gamma element features on speed could be assessed by the redundancy between discretized gamma element features *f* and the speed variable *V*, i.e. *Red*(*f*;*V*) = *I*(*f*; *V*). Such redundancy was then normalized by the entropy of *f* to quantify the fraction of information about the variability of *f* explained by the variability of *V* (cf. **Fig. 3f**).

### Statistical analysis

All statistics were performed using either built-in Matlab (R2021a) functions, Matlab toolboxes, or Statistica 13.

Electrophysiological data was analyzed on all 40 trials per animal except for mice mouse #3 (missing trial 15) and mouse# 4 (missing trials 21-24) due to technical issues with the electrophysiological recording.

#### Behavior

Average latency to reach the reward during the learning phase of the task (i.e., days 1-10) was analysed by a non-parametric Friedman ANOVA (within-factor: days). For the probe test, we carried out a repeated measure ANOVA (within-factor: arms) on the ratio of number of visits in each arm compared to chance level (0.125). Post-hoc tests were used when appropriate. For all analyses, the significance threshold was 0.05.

#### Distributions of gamma bouts features

The probability density function (pdf) of each gamma feature (amplitude, frequency and theta-phase) was established across all trials, per electrode. As the pdfs of gamma frequency often displayed a wide range whatever the channel and the animal, we restricted most of our analyses to one representative channel per anatomical layer: the channel displaying the pdf the most consistent with expectations from the dual-gamma band literature, usually based only on very strong gamma episodes (here: the strongest 5 % gamma bouts). Hence, we favored channels showing a dominance of lower gamma frequencies in the *rad*, and faster gamma frequencies in the *l-m*. The mean pdf of each layer was calculated across mice, before generating an artificial sample of the relevant gamma feature matching this pdf (n=10000; random sample with replacement). For the frequency, the statistical difference in the distribution from each pair of layers was evaluated using a bootstrap method (2000 repetitions using 1500 subsamples) on the Kullback-Leibler divergence^52^ so that to be significant, the mean divergence between separate layers had to be greater than the upper 95% confidence interval on the mean divergence between shuffled layers. In addition, both the modes of the individual mouse frequency pdfs and the ratio between gamma_M_ and gamma_S_ were compared across layers using one-way ANOVA or non-parametric Kruskal-Wallis tests on the between-factor ‘layer’, with post-hoc comparisons. The ratio was defined as:

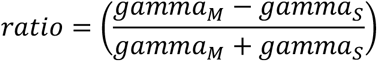

with gamma_S_ and gamma_M_ being the sum of probabilities for the individual mouse frequency range centered on the frequency modes between 25-50 Hz and 60-100 Hz, respectively, and whose probability is ≥ 50 % of this mode.

For the phase, all statistical analyses were carried out using the ‘circStat’ Matlab toolbox^53^. First, the statistical difference between artificial distributions from each pair of layers, generated as for the frequency mean pdf, was assessed using the Kuiper two-sample test (note that very similar results were obtained using the circular Watson’s U2 test with 1000 permutations). Second, their mean phases were compared by pairs of layers using the Watson-Williams test for circular means after checking the non-uniformity of these distributions (omnibus test). This latter analysis was also done on the individual mouse distributions (n=5) to compare the grand mean phase across layers. All the above analyses were performed iteratively on distributions containing a varying range of data, from all data (0^th^ percentile: no further data selection) to the 95^th^ percentile of the maximum amplitude (i.e., only the gamma bouts with the 5 % strongest amplitude), by steps of five percentiles. Percentiles were calculated for each trial and electrode before pooling the bouts from all trials per electrode.

#### Classification, cross-classification and mutual information

For single mouse performance levels (fraction of classification correct and confusion matrices), as well as for information theoretical quantities we evaluated 95% confidence intervals using a bootstrap with replacement approach (1000 replicas) over the lists of gamma elements retained for inclusion in each of the analyses. When comparing multi-mouse samples of performance metrics or testing their significance against a threshold (as in the boxplots of **Fig. 3-4** and **Extended Data Fig. 9**), we used t-test (two-tailed for inter-sample comparisons and one-tailed for comparison of single samples against a chance-level or zero threshold). We report uncorrected p-values in captions and text, however significance, unless specified otherwise, is assessed using Bonferroni correction for multiple comparisons (*, **, *** denote corrected p-values smaller, respectively, than 0.05, 0.01, 0.001; symbols in brackets indicate significance only prior to multiple comparisons correction; when significant deviations in both directions above or below chance level occur, we use upward ↑ or downward ↓ symbols instead of *’s). Boxes in the boxplot mark the inter-quartile range (IQR), the horizontal line sample mean μ, the whiskers μ ± 2*σ where σ is sample standard deviation.

## Code availability

Available upon reasonable request.

## Acknowledgements

This work was supported by CNRS, INSERM, Université de Strasbourg, Aix-Marseille Université and grants from the University of Strasbourg Institute for Advanced Studies (USIAS grant to DB) and from the Agence Nationale de la Recherche: ANR ERMUNDY (ANR-18-CE37-0014) to DB, MdV and AT; ANR DG-Goal (ANR-17-CE37-0002) to RG; ANR HippoComp (ANR-21-CE37-0011) to RG and DB, Labex MME-DII (ANR-11-LBX-0023-01) to AT. The authors wish to thank Pr. Jesse Jackson for critical reading of the manuscript and the Buzsaki’s lab for publicly sharing their data.

## Author contributions

VD and RG performed the experiments and analyzed the data, MDV and AT built the model, DB analyzed the data and built the classifier. RG and DB conceived the experimental and analytical design; all authors wrote the article.

## Competing interests

The authors declare no competing interests.

## Materials & Correspondence

Demian Battaglia and Romain Goutagny

## Notes

### Competing Interest Statement

The authors have declared no competing interest.

