## Supplementary figures and captions for "Hippocampal gamma oscillations form complex ensembles modulated by behavior and learning"

### Extended data

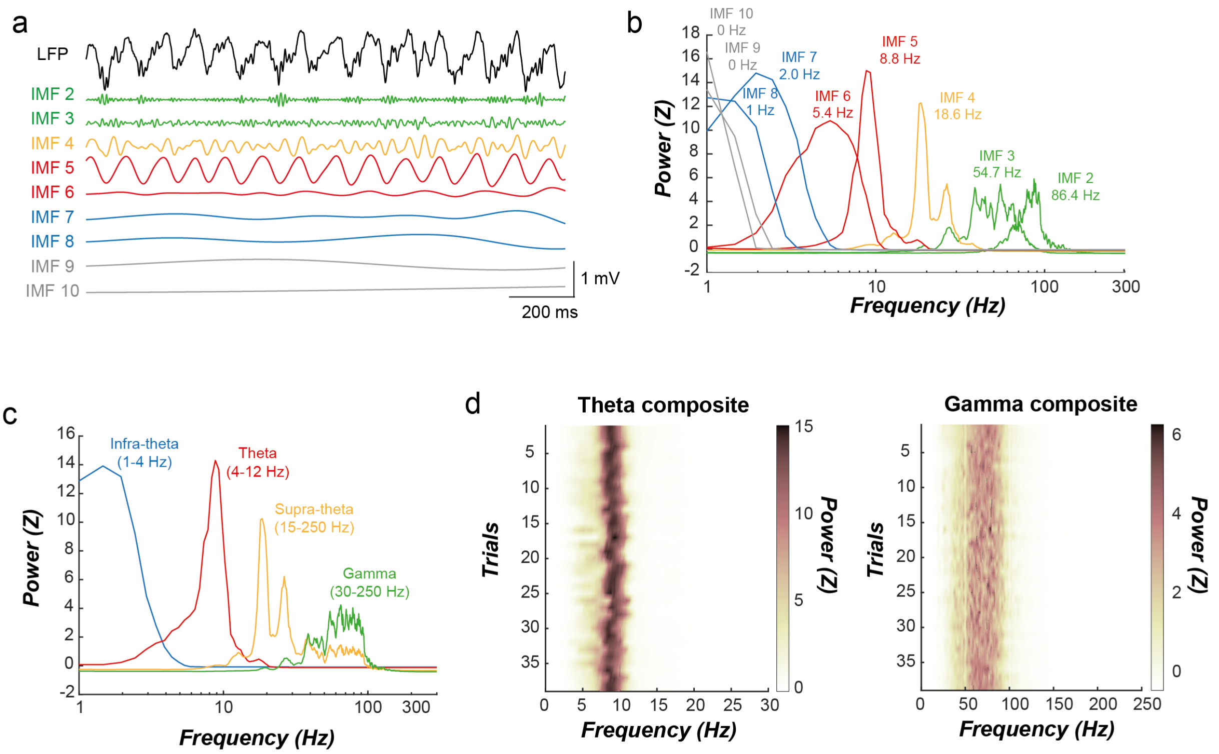


**Extended Data Fig 1: EEMD decomposition of recorded LFPs and CSDs.**

**a,** Each LFP is decomposed by EEMD, independently by trial and channel, into 10 IMFs. A portion of the LFP and derived IMFs traces is shown (IMF 1 corresponds to the residual and is ignored further). Each IMF is color-coded depending on the frequency band in which its mean instantaneous frequency falls in (infra-theta: 1-4 Hz; theta: 4-12 Hz; supra-theta: 15-250 Hz; gamma: 30-250 Hz). **b,** Z-scored power spectrum of each IMF with its peak frequency. **c,** Z-scored power spectra of the composite signals for each frequency band. **d**, Stability of the Z-scored spectral contents of both theta (left) and gamma (right) composites of a channel across trials. Examples from mouse #3.

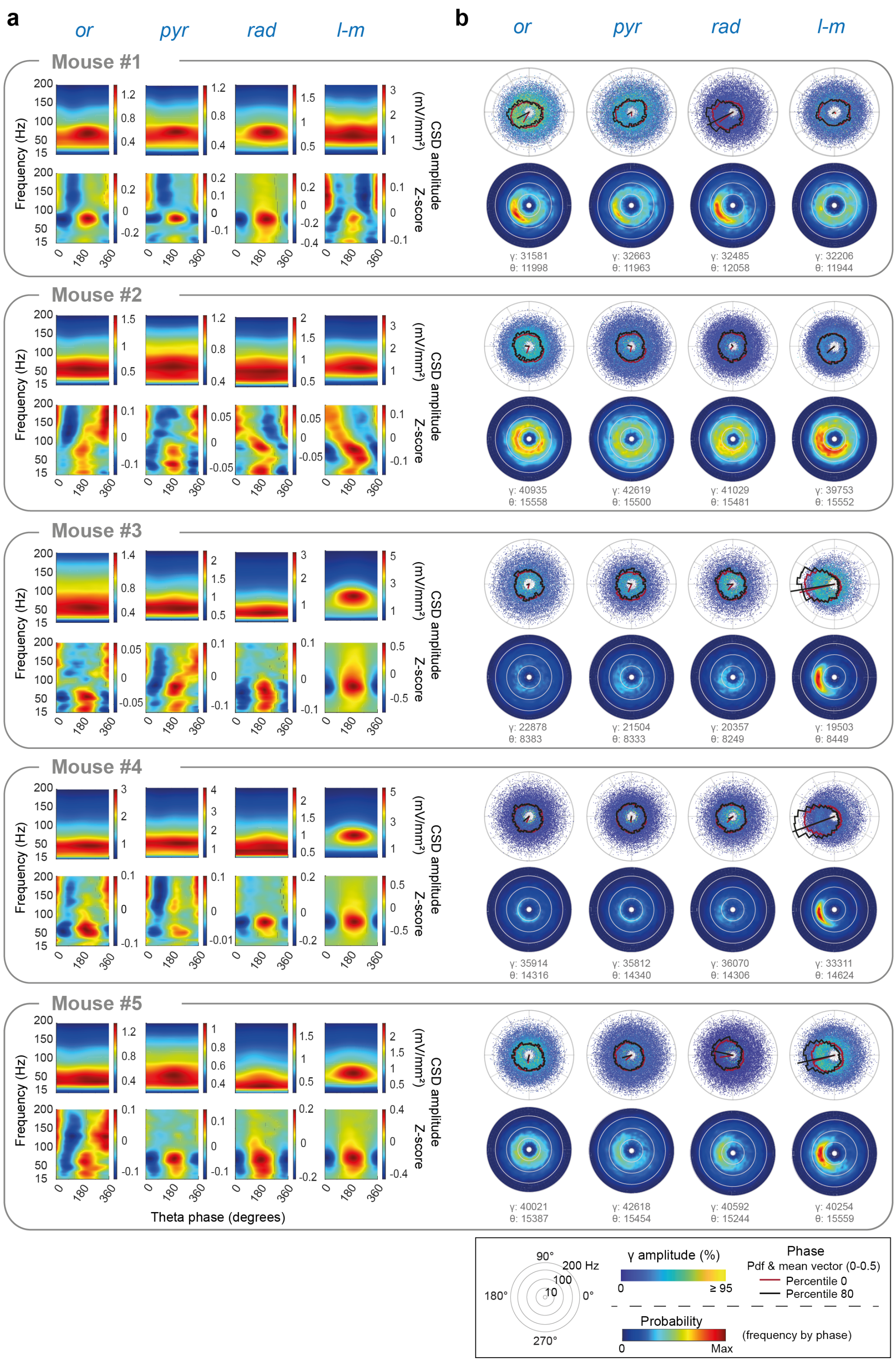

**Extended Data Fig 2. Consistency of the gamma diversity across layers and mice.**
**a**, The average gamma amplitude per theta-phase (top row: raw; bottom row: z-scored across phases) indicates that the *l-m* has a higher average gamma frequency content, especially compared to the *rad* in accordance with the literature (at the exception of mouse #1 which present gamma$_M$ in both *rad* and *l-m*) **b**, Top row: Nonetheless, the distributions of theta-nested gamma elements show that, across mice and layers, these transient gamma bouts can occur at all frequencies, amplitudes and theta phases, even if the *l-m* has more elements concentrated towards the theta trough (3 out of 5 mice). Theta phase pdf estimates and mean vectors are overlaid for either all gamma bouts (percentile 0; red) or a subset of only

high amplitude gamma bouts (percentile 80; black). Bottom row: The probability of occurrence for gamma elements per frequency bins (10-Hz bin) for each phase (10° bin) shows that, independently of the gamma amplitude, the gamma elements in the *l-m* tends to occur preferentially in $gamma_M$ around theta trough, in line with previous studies. However, a relatively similar pattern can be observed, although to a lesser extent, in other layers, strengthening the claim that average-based analysis is biased towards the high amplitude elements. Theta cycle and gamma element counts for each representative channel over the ATA learning are indicated underneath.

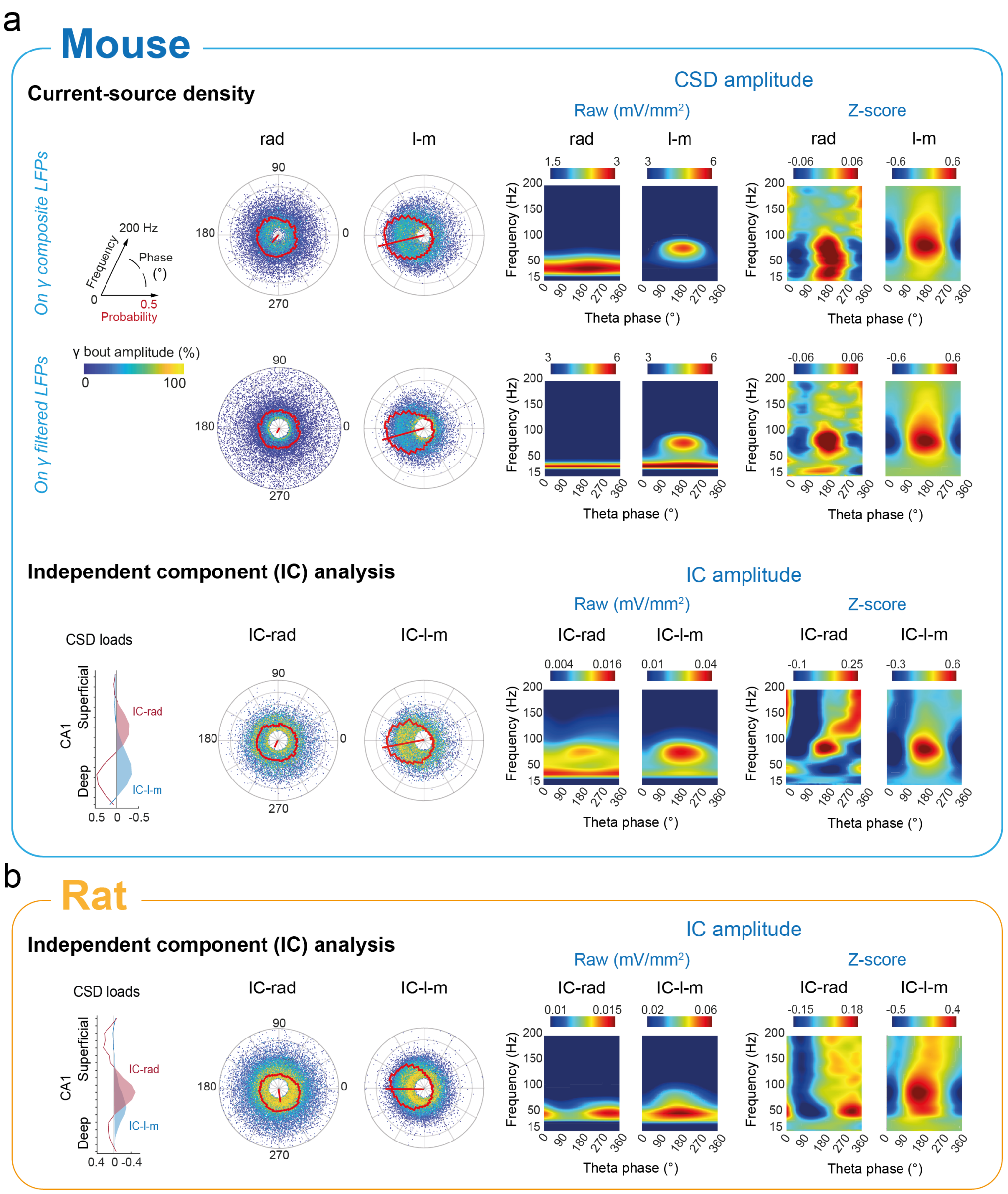


**Extended Data Fig 3. Theta-nested gamma diversity does not depend on processing methods or rodent species.**

**a**, top row: Gamma elements present an overall similar diversity of features (left; red overlay: pdf and mean vector for theta-phase) whether they are extracted from EEMD-derived or filtered (zero-phase filtering using a finite impulse response filter) gamma signal, with the theta-phase being determined from the EEMD-derived theta signal using either Hilbert transform or waveform-based linear interpolation between cycle quadrants [1]. Of note, the EEMD method seems less susceptible to potential theta harmonics around 30 Hz as evident in the average theta-gamma coupling motifs (right). Example from mouse #3 (percentile 0; one representative channel for *rad* and *l-m*). Bottom row: Broadly similar gamma diversity was obtained using independent component analysis, a decomposition method aiming at separating different spatio-temporal components within a multi-channel mixed signal (fast kernel density independent component analysis [ICA] algorithm from the ‘ICAofLFPs’ Matlab toolbox; [2]), instead of our individual channel decomposition approach (a). Applied on all CA1 LFP channels (same dataset as above), filtered beforehand in the gamma band (zero-phase filtering using a finite impulse response filter with 30-250 Hz passband), two independent components could be assigned respectively to str. *rad* and str. *l-m* based on their voltage (V) and their second spatial derivative (CSD) loadings along the channels (left). Note that theta cycles for the ICA method were extracted on filtered theta. **b**, Gamma features diversity is also present in rat CA1 recordings during locomotion (concatenated epochs of high theta power during a linear track task [3]). Example shown is from rat AB3 (session 60, shank 5). Applying our individual gamma bout extraction method to fast kernel density ICA-derived components from CA1 gamma-filtered LFPs reveals that gamma elements occur at various frequencies and (filtered) theta-phases in both components. The theta-phase difference between the respective average theta-gamma spectrograms might however be stronger in rats than mice.

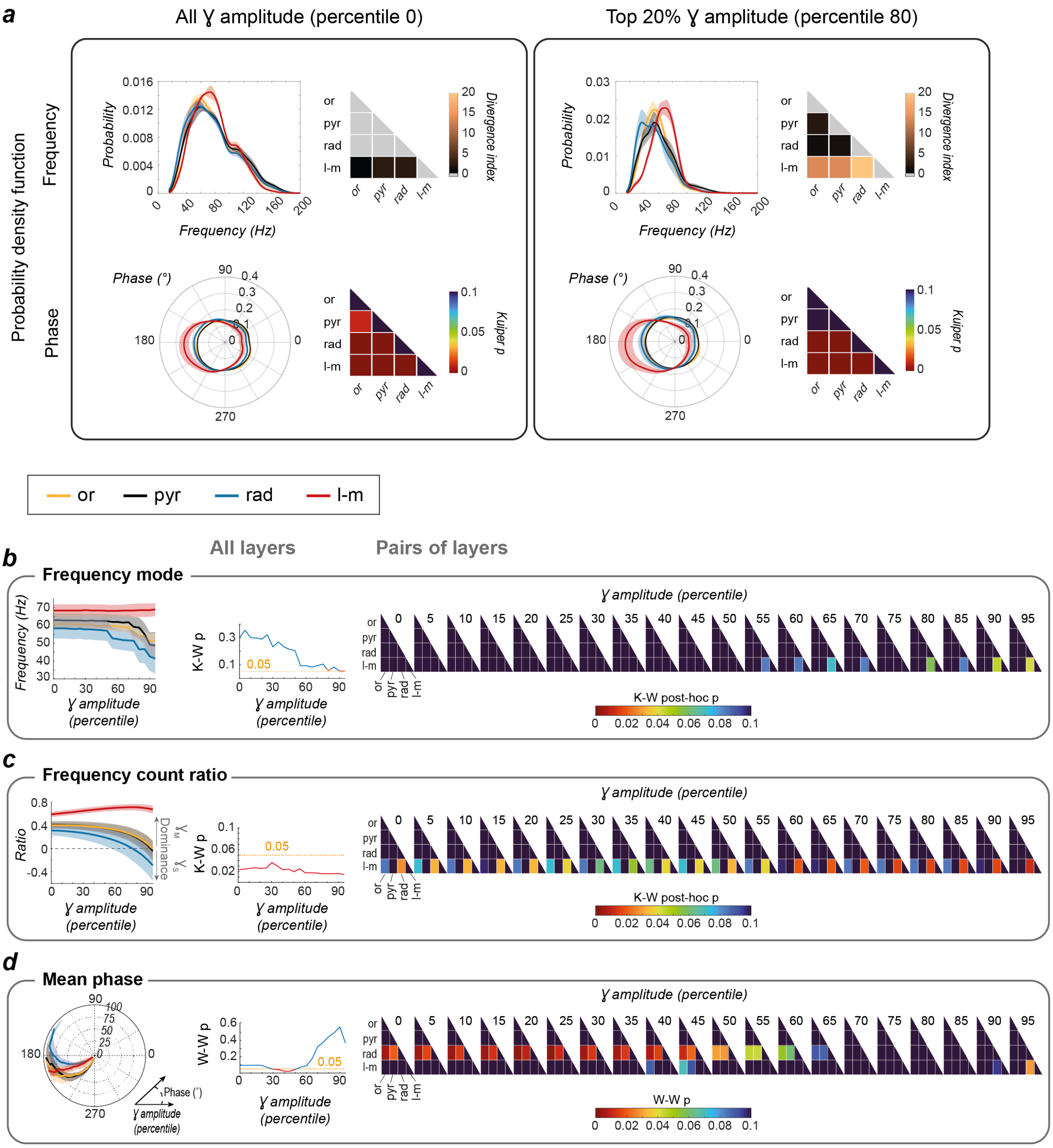


**Extended Data Fig 4. Gamma elements in different layers have different continuous distributions of frequency and phase.**

**a,** *Top row:* The difference in gamma frequency contents of CA1 layers increases with the restriction to stronger bouts, especially between the *rad* and the *l-m*. First, distributions of gamma bouts frequencies of each pair of layers were generated based on the mean pdf (± SEM; n=5 mice; 10000 samples) from each CA1 layer for either all their gamma elements (i.e., percentile 0) or only those in the top 20% gamma amplitude (i.e., percentile 80) recorded during the ten days of learning. Their distributions within a percentile were then compared, by pairs of layers, using a bootstrap method on the Kullback-Leibler (KL) divergence, a measure of the distance between two distributions. The more the divergence index differs from zero (significance threshold at alpha = 0.05), the more significant the difference (i.e., any non-grey part of the KL divergence matrix is significant; values were rounded to the nearest integer). Overall, the *l-m* always differs significantly from the *rad*. Considering only the strongest gamma elements exacerbates this difference. *Bottom row:* The theta phase distribution differs between CA1 layers, with a stronger phase-locking around theta trough in the *l-m*. Although the other three layers are visually closer to a uniform distribution (but omnibus test p values < 0.01), only the *or* and the *pyr* do not statistically differ for some gamma amplitude percentiles (Kuiper test for circular distributions comparing pairs of samples generated from the mean pdf; n=5 mice; 10000 sample size). Mean pdfs (± SEM) and p values of their comparison by pairs are shown for gamma amplitude percentiles 0 and 80 but each layer phase distribution remains quite similar across percentiles. **b,** *Left:* While the frequency mode for the *l-m* remains constant (peak of the pdf: ~ 70 Hz) across percentiles, the modes from the superficial layers become lower with increasing percentiles although a general difference almost reached significance only for some very high amplitude percentiles (*center:* Kruskal-Wallis [K-W] test p values). *Right:* Comparing pairs of layers revealed that only the *rad* and *l-m* differed significantly (matrices of K-W p values ceiled at 0.1) from percentile 55. **c,** *Left:* The dominance of $gamma_M$ over $gamma_S$ frequencies (ratio of probability sum in 25-50 Hz and 60-120 Hz ranges respectively) is inverting with high amplitude percentiles for all layers but the *l-m*, with significant difference between layers from percentile 30 (*center:* K-W test p values). *Right:* Comparing pairs of layers confirms that the *l-m* always contains more $gamma_M$ than the *rad*, significantly from percentile 30 (*right:* matrices of K-W p values ceiled at 0.1). **d,** *Left:* The grand mean angle (i.e. circular mean of individual mice's preferred theta phase) tends to become earlier as the amplitude percentile of gamma bouts increases, except for the *l-m*. A significant difference between layers is present for percentiles 30 to 50 (*center:* Watson-Williams [W-W] test p values). *Right:* Comparing pairs of layers suggests mainly that the *rad* differs from the *or* and the *pyr* at percentiles 0-55. It is however worth noting that the distributions of the individual mean phases were not different from uniformity (omnibus tests: p = 0.31) although the Rayleigh test, a test with more assumptions regarding the data distribution, indicates significant (or close: p ≤ 0.06) p values for all conditions except for percentiles 90-95 of *str. or*.

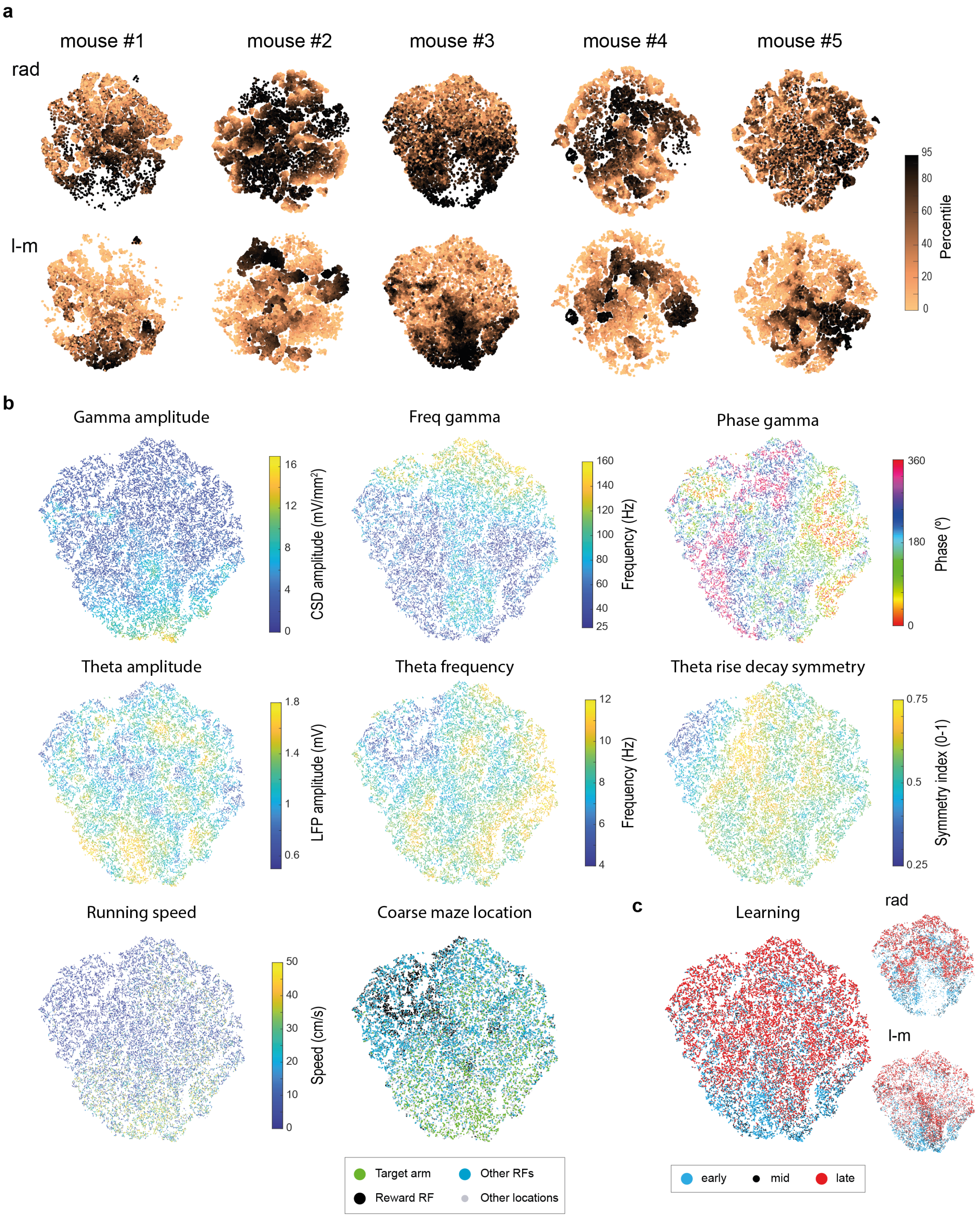


**Extended Data Fig 5. Diversity of gamma elements from both CA1 distal dendritic layers largely overlaps and cannot be explained completely by simple behavioral variables.**

**a,** Distributions of *rad* and *l-m* gamma element ensembles largely overlap in all mice. When plotting, separately for the *rad* and *l-m*, the results of the t-SNE calculated on their pooled gamma elements features, it appears that both layers display very similar distribution patterns. Some exclusive territories exist but are constituted, in most cases, of elements with a large gamma amplitude. Note that the *rad* seems to have a more varied combination of features for strong gamma elements than the *l-m*. **b-c,** The representation of the running speed, coarse maze location or learning stage on matching dimensionally reduced representations suggest that these behavioral variables would entertain complex relationships with combined features. *Example from mouse #3 (all gamma elements from the 10-day ATA learning; rad and l-m representative channels).*

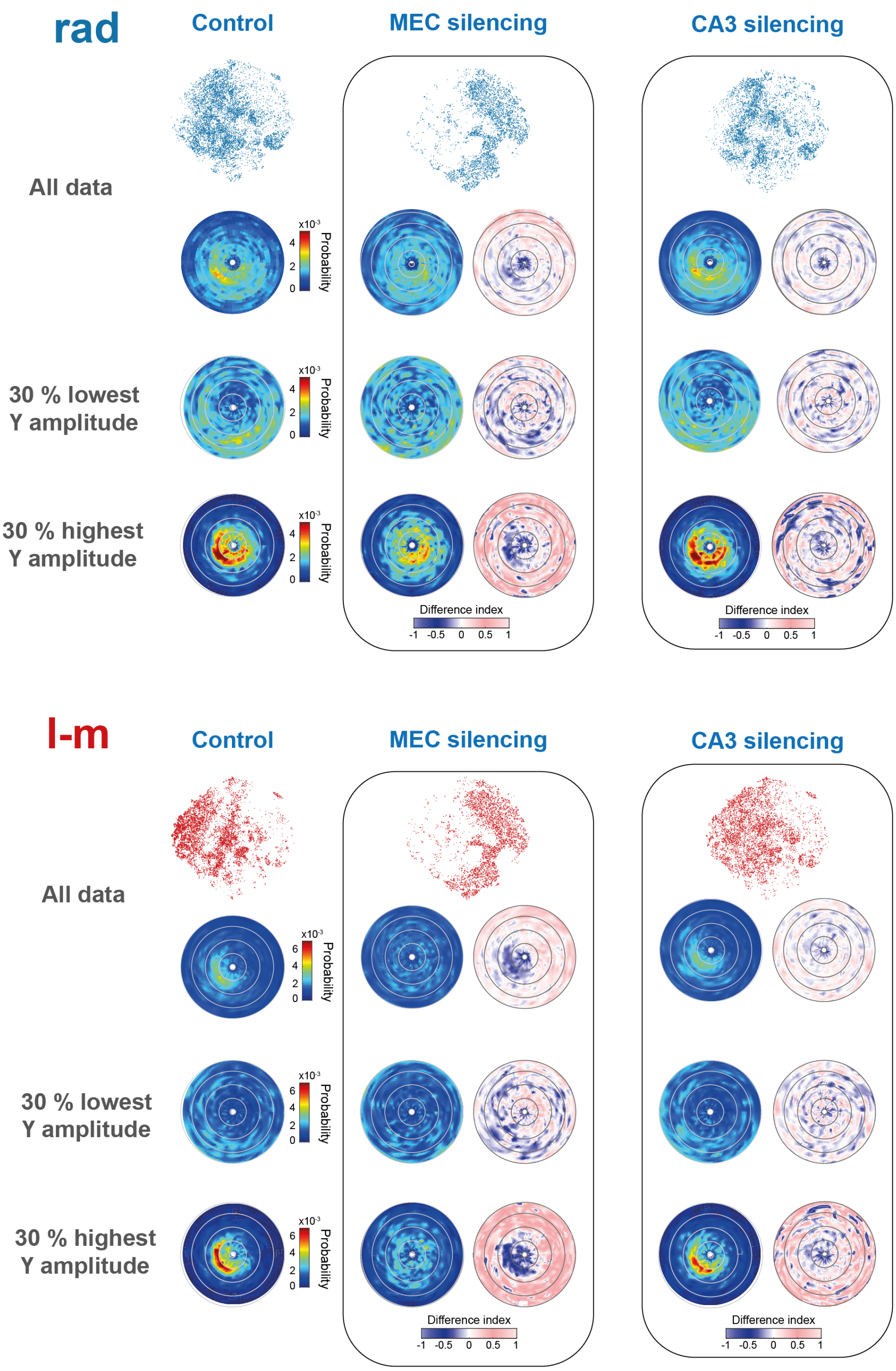


**Extended Data Fig 6. Effect of medial entorhinal cortex (MEC) or CA3 inactivation on gamma elements diversity in the rad and l-m layer.**

*Top row*: t-SNE dimensionality reduction made with the 6 features gamma elements vectors. Inactivation of either the MEC or CA3 change the gamma landscape in both the *rad* and *l-m*. Note however that MEC inactivation drastically impact gamma elements landscape in both layers.

*Below the t-SNE*: the probability of occurrence for gamma elements per frequency bins (10-Hz bin) for each phase (10° bin) and the difference index maps show that inactivation of the medial entorhinal cortex (MEC) or CA3 removes gamma elements at all phases and frequencies. Importantly tough, the loss of frequency-phase preference in the *l-m* following MEC inactivation is mostly driven by the loss of high gamma amplitude elements (top 30%) given that no such preference is observed when considering only the weakest elements (30% lowest gamma amplitude). Further, both manipulations impacted all CA1

sub-layers and neither of them suppressed the diversity of gamma elements, showing that inputs do not elicit gamma in separate, narrow frequency and phase bands, but convey complex gamma ensembles (with frequency differences evident only for high power events).
Examples from mice IZ33 and IZ27 as in [4].

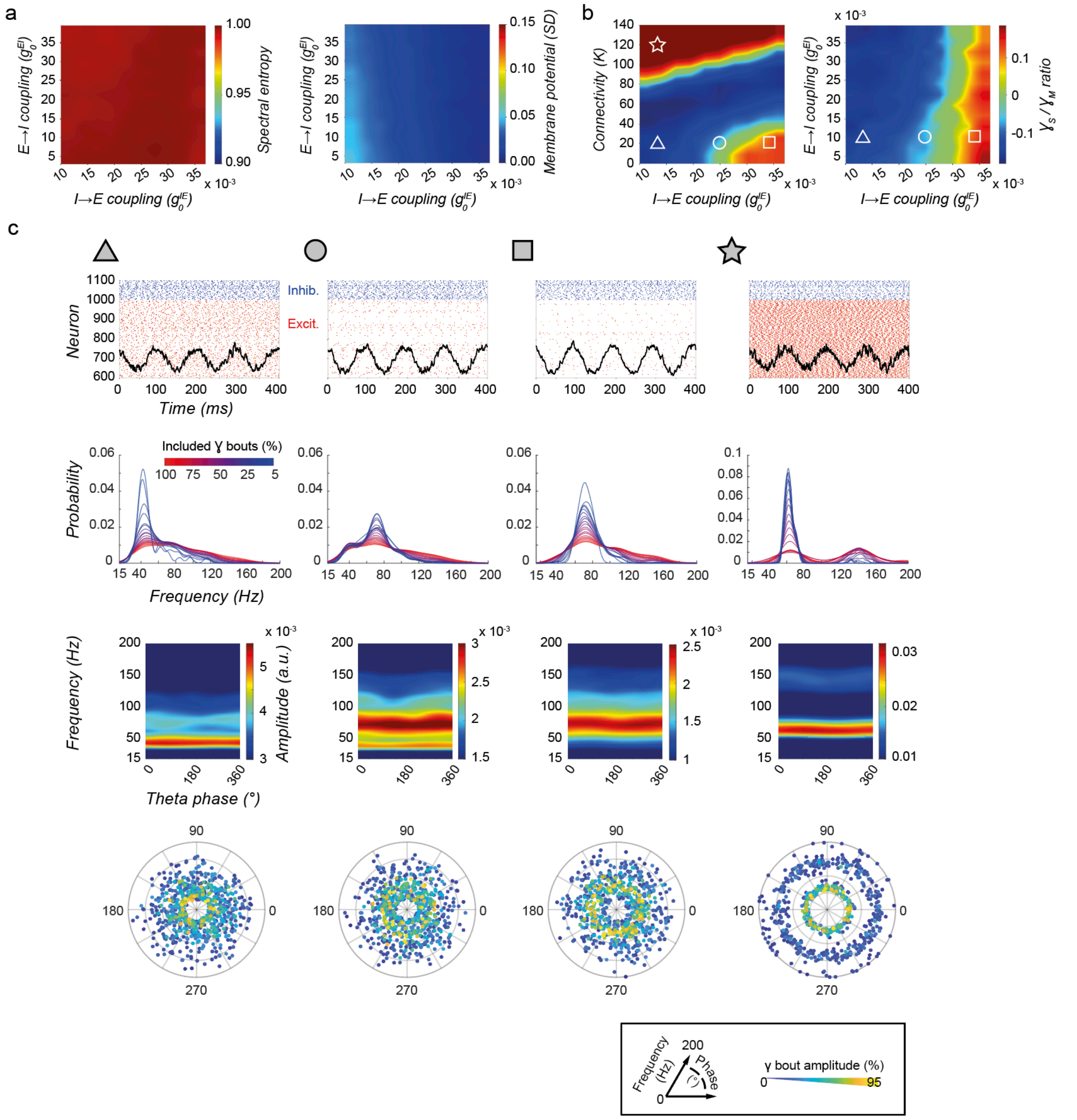


**Extended data Figure 7. Additional details on computational model.**

**a**, Surfaces of parameter dependency of spectral entropy of the LFP-like signal and the standard deviation of the mean membrane potential of inhibitory cells, as in Fig. 2e but here as a function of the I-to-E and of the E-to-I recurrent synaptic connectivity conductance strengths. Both quantities display a weak dependency on changes of these parameters, i.e. the fringe-of-synchrony regime is very robust, as long as global connectivity density remains moderate (cf. Fig. 2e, right, and star working point). **b**, The ratio between gamma elements with frequencies in a slow and in a medium frequency range can be modulated by adjusting parameters. When remaining in a fringe-of-synchrony regime (triangle, circle and square working points, low connectivity density), increasing the strength of the I-to-E synaptic conductance increases the fraction of gamma elements with slow-range frequencies. For high connectivity density, in the highly synchronous oscillations regime (star working point), frequency ratio is instead mostly dominated by slow frequency power. **c**, Nature and properties of the different considered working points for different parameter choices. From top to bottom: representative raster plots of simulated spiking activity; normalized counts of detected gamma elements for different frequencies, filtered to include elements with gamma amplitude above different quantiles (red, all elements; dark blue, top 25% of amplitudes only); average single theta-cycle spectrograms of LFP-like signal; polar scatter plot representation of different elements (as in Fig. 1d and 2d). The shift from faster to slower frequencies in both the spectrograms and the peaks of elements counts distributions (associated to high amplitude events) is evident when moving from the triangle to the square, via the circle working points. For the unrealistically synchronous star working point, the events have frequencies concentrated within narrow frequency peaks only. For all the other working points, gamma element diversity in frequency is pervasive. The model does not capture the phase-selectivity of strong amplitude gamma elements displayed by the data (see horizontal stripes in single-theta cycle spectrograms, denoting lack of phase concentration).

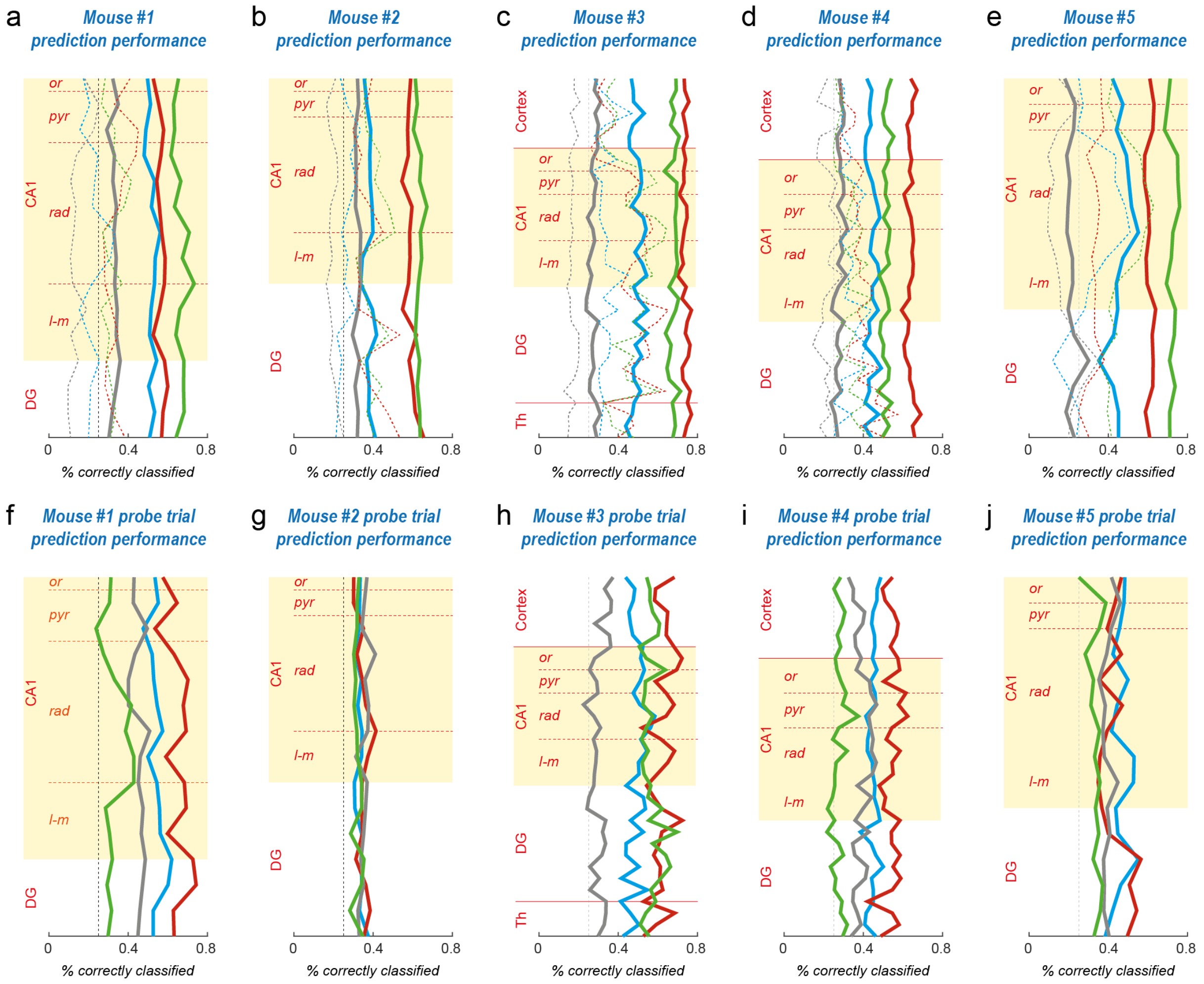


**Extended data Figure 8. Maze location decoding performance for all mice and for alternative classifiers or trials.**

**a-e**, As in Figure 3c for mouse #3 but here for all mice, we show classifier decoding performance (fraction of correctly classified locations), by maze location (colors as in Figure 3b: red reward field, green target arm, light blue other end-arm fields, grey other locations). Different classifiers were trained for different depths along the dorsal hippocampal axis. Solid lines indicate average performance across all trials. Performance in detecting target arm, reward field and other arm end fields was significantly above chance level for every anatomical layer and all mice, and, for mice #1 and #2, performance in detecting other locations as well (comparison of 95% bootstrap c.i. with chance level, dashed black line). Confidence interval, not shown to avoid figure overloading, were narrow, similarly to Figure 3c, and bounded by mean ± 0.07. Dashed lines indicate performance of classifiers trained on gamma features only (cf. Figure 3e, right). **f-j,** Performance of same classifiers as in panels **a-e,** in detecting different maze sections based on elements measured in probe trials. Confidence interval, not shown to avoid figure overloading, bounded by mean ± 0.12. For all mice, but mouse #3, classification in decoding target arm approached chance level (and became not significant for most channels in mouse #4, panel i).

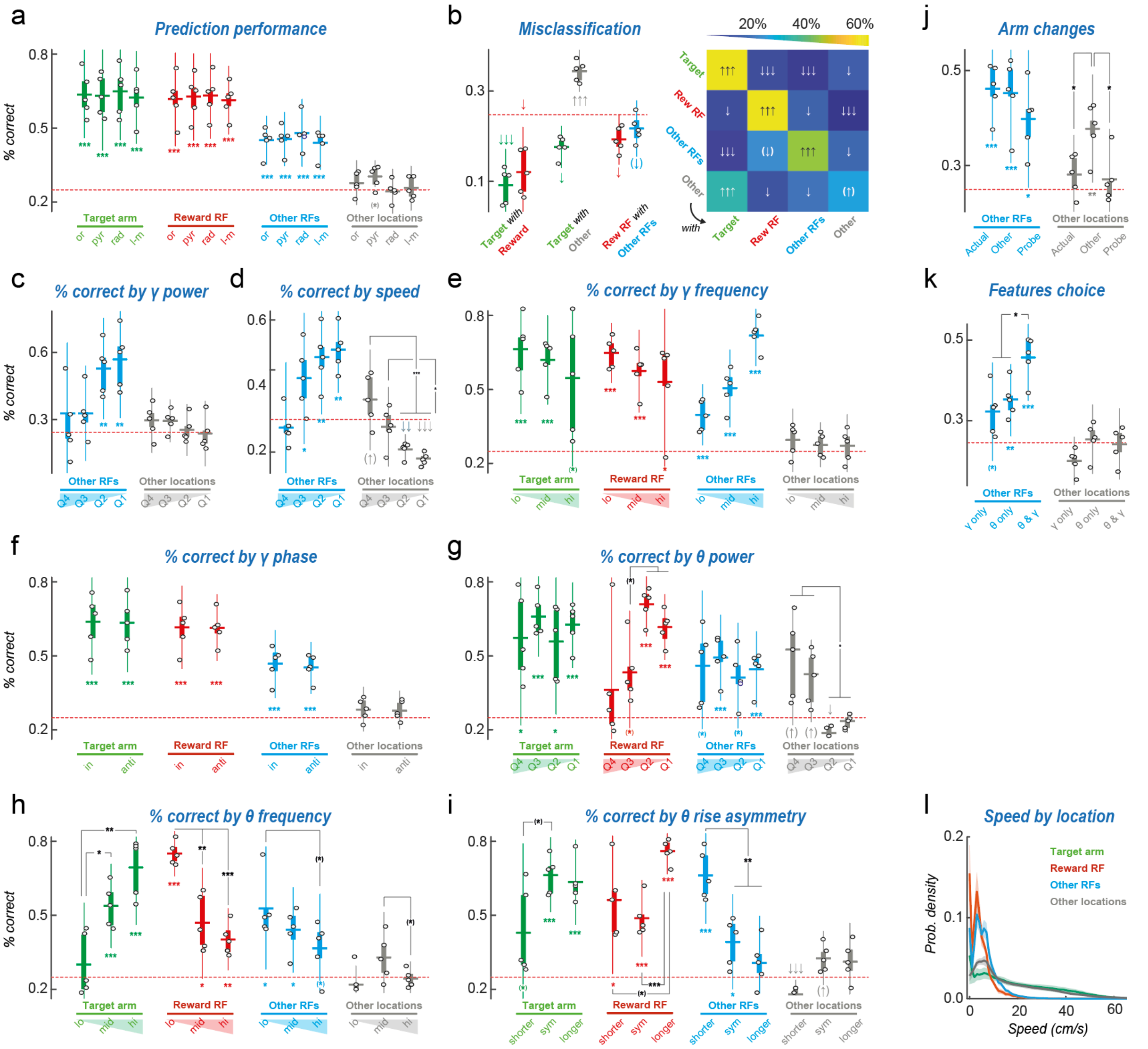


**Extended data Figure 9. Additional information about factors affecting decoding performance.** In boxplots, dots represent performances of individual mice; box, IQR; horizontal line, sample mean; whisher, 95% sample c.i. One-tailed t-test for comparison of sample vs chance level (colored symbols for significance); two-tailed t-test for comparisons between samples (black symbols). *, $p < 0.05$; **, $p < 0.01$; ***, $p < 0.001$ after Bonferroni correction; symbols in brackets denote significance only before Bonferroni correction. **a,** Performance in decoding different maze locations averaged over gamma elements and trials for different CA1 anatomical layers. Performance depends on maze location class, but is stable across CA1 layers. **b,** Analysis of misclassification patterns. Confusion matrix to the right (averaged over mice and trials; significance of entries, ↑↑↑, ↑↑, ↑ above chance level; ↓↓↓, ↓↓, ↓ below chance level, with p < 0.001, 0.01, 0.05). To the left, detailed boxplots for some types of confusion pattern to the left. Among the confusions occurring at less than chance level, denoting classes that are very efficiently discriminated: target arm and reward field ($p < 0.0009$ for target arm classified as reward and $p < 0.002$ for reward classified as target arm); target arm with other locations ($p < 0.002$ for target arm classified as other location); or yet, reward field with other end-arm fields ($p < 0.009$). However, other locations were confused between them only barely below chance level (generic box decoded as reward box, $p < 0.0312$, not significant after Bonferroni correction) or even above chance level denoting thus a tendency to systematic error (other location decoded as target arm, $p < 0.001$). **c-i**, Performance in decoding location by different locations as a function of the values of specific features of gamma elements. **c-d**, Performances by quartile of gamma amplitude (**c**) and quartile of motion speed (**d**) in decoding other end-arm fields and other locations (see Fig. 3d for target arm and reward box).

Remarkably, the “Other location” zone, usually difficult to discriminate above chance level, can be significantly decoded if speed is large (e.g. $p < 0.016$ prior to Bonferroni correction, for top quartile of speed, i.e. the same quartile for which decoding of target arm is also the most efficient, cf. Fig. 3d, so that a proper discrimination between generic and target arm is possible if speed is not too slow). **e-f**, performance as a function of gamma element frequency (**e,** low, medium or high frequency ranges) or phase (**f,** around 0° or around 180° theta phases). **g-i,** performance by quartiles of theta amplitude (**g**), frequency (**h**) and cycle waveform asymmetry (**i**). **k-l**, performance in decoding other end-arm fields and other locations: **k,** when training classifiers to decode a generic alternative arm (“Random”) or when using standard classifiers but applied to probe trials (“probe”); or, **i**, when training classifiers to use only gamma or only theta input features. See Fig. 3e for performances in decoding target arm and reward field locations. **l**, to further assess the role of speed in allowing decoding, we studied how the distributions of speed varied by the different maze locations. Distributions for the reward field (RF) and the other end-arm fields (other RFs) were peaked on slower speeds, while distributions for target and other arms had a longer right tail toward larger speeds, associated to faster exploration movement. Note that the highest confusions in classification arise between locations with similar speed distribution types (cf. panel **b**). Nevertheless, all the locations display a large variety of speeds, and there is substantial overlap between the distributions for all locations. Therefore, the classifiers, to achieve successful location decoding must extract from gamma elements some additional information besides an inference of current speed.

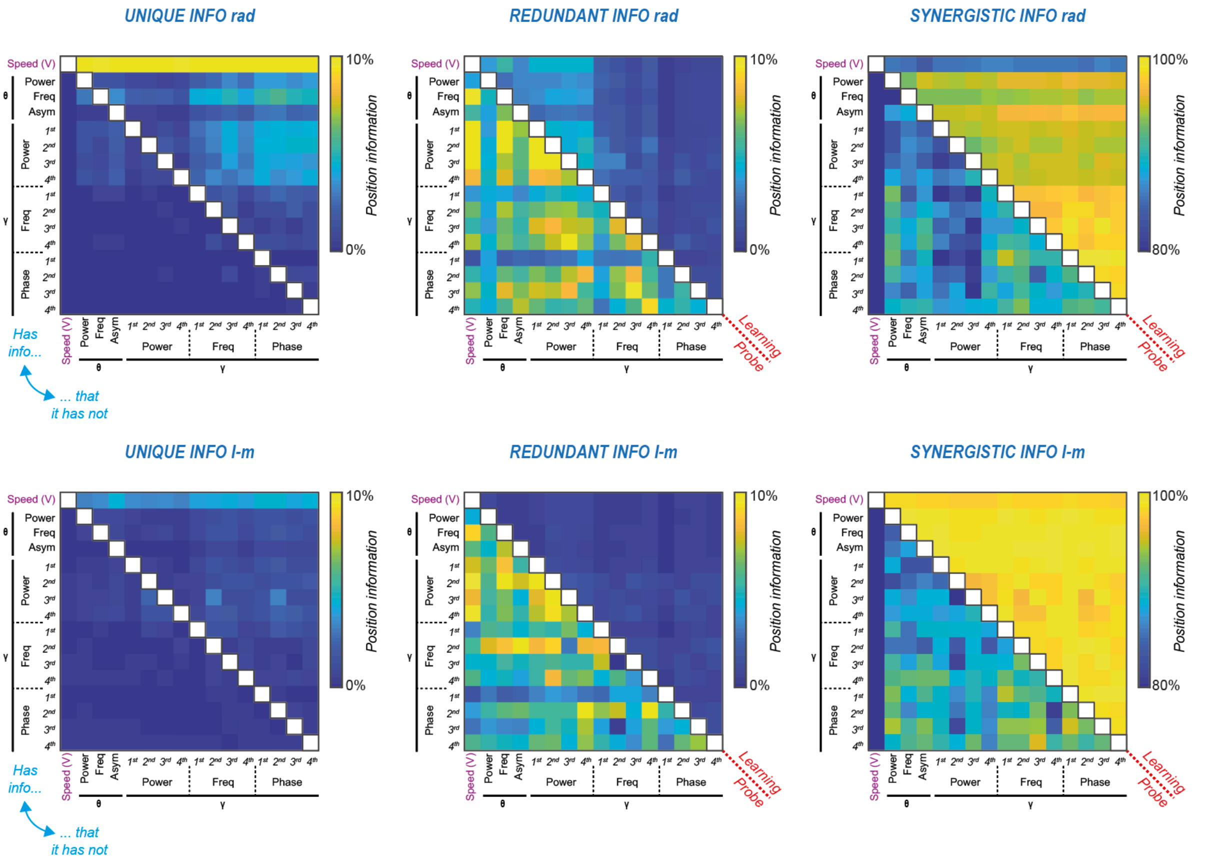


**Extended data Figure 10. Additional details about information theoretical analyses.** Decomposition of mutual information between input feature pairs (the rows and column of the shown matrices) and maze location, into unique (left), redundant (middle) and synergistic (right) fractions, averaged over all mice. Top row, pairs of features from a representative *rad* channel; bottom, from a representative *l-m* channel. The two triangular parts of the matrices describe, respectively, information decompositions for: learning trials (upper triangular part) and probe trials (lower triangular part). For all considered pairs of features, most of the mutual information with maze location is of synergistic nature (note the different color bars for the three different columns). In learning trials, synergistic information fractions are larger for *l-m* than for *rad* pairs of features (two-tailed t-test comparison between pairs of lists of triangular part entries, $p < 0.0001$) while redundant and unique information fractions are larger for *rad* than *l-m* ($p < 0.0001$), particularly the unique information conveyed by speed but none other gamma element features ($p < 0.0001$). This finding agrees with the hypothesis that *rad* activity reflect inferences based on an internal CA3 model. It should thus be less modulated than *l-m* activity (reflecting encoding) by the actual speed of ongoing navigation behavior, so that the unique information about maze location conveyed by speed should be larger. On the contrary, *l-m* gamma elements can support an encoding of maze location in a way synergistic with speed, as revealed by larger synergistic fractions in *l-m* than in *rad* for feature pairs including speed ($p < 0.0001$). In probe trials, synergistic fractions are decreased and redundant fractions increased for both *rad* and *l-m* ($p < 0.0001$), denoting altered statistics of gamma element production in probe trials and “broken” synergistic encoding of location.

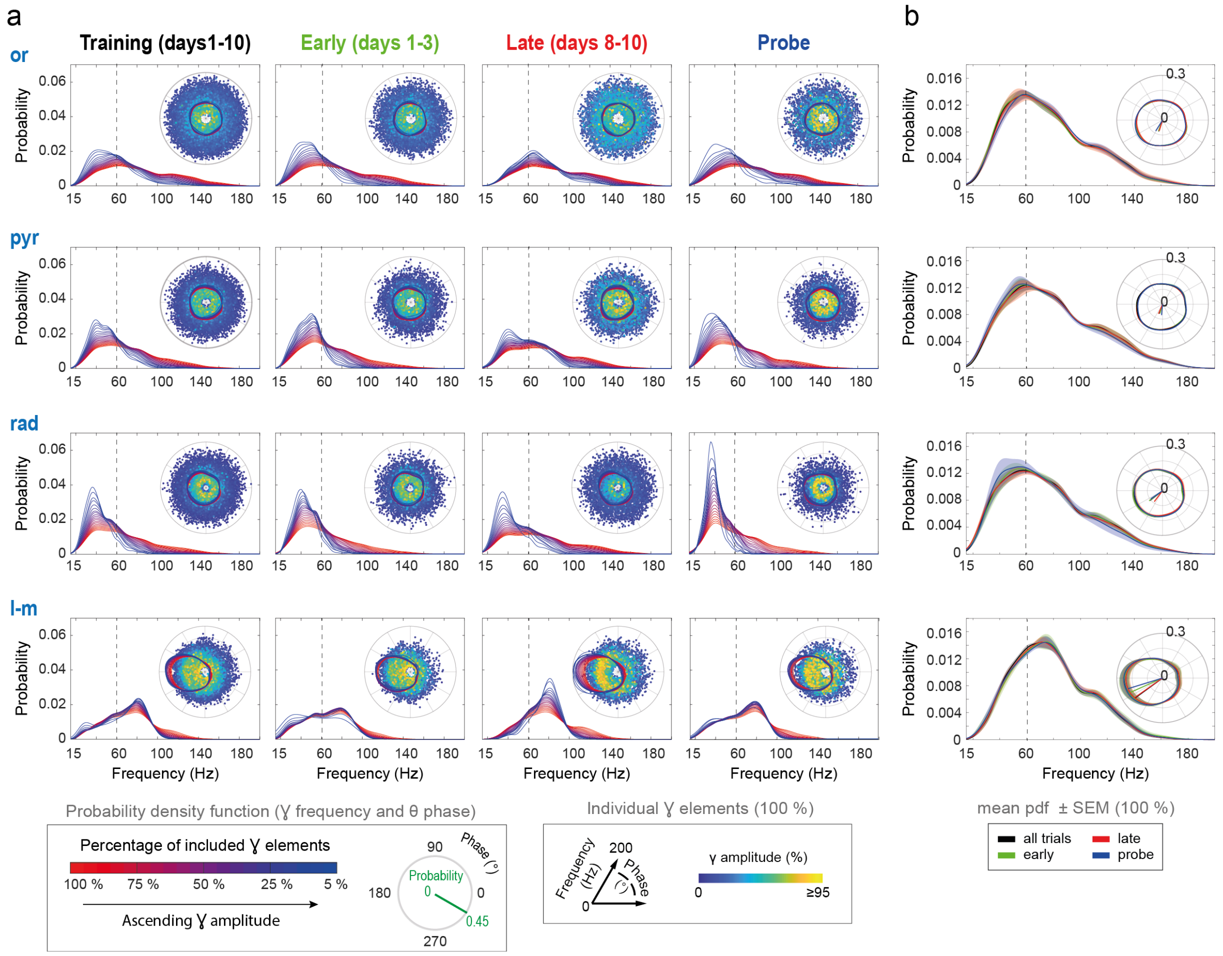

**Extended data Figure 11. Frequency and phase of hippocampal gamma elements are continuously distributed at all learning stages.**

**a**, Example (mouse #3) of pdf estimates of the gamma frequency for the gamma elements recorded on the representative channel of each CA1 layer, color-coded by the amount of data included (from 100% in red to the most conservative top 5% gamma amplitude in blue), for different periods of the task: all training, early, late and probe trials. Inserts: joint distribution of the three gamma features (amplitude, frequency and theta-phase) for all gamma elements (i.e., percentile 0: 100% of data) with overlaid pdfs for the theta-phase (color-coded by gamma amplitude percentile). No dramatic change can be observed in the occurrence of either slower versus faster gamma frequencies or the theta-phase, whichever the task periods. The large diversity in the gamma element features thus remains rather stable across learning. **b**, Frequency and phase distribution across learning and anatomical layers in all mice (± SEM; n=5 mice).

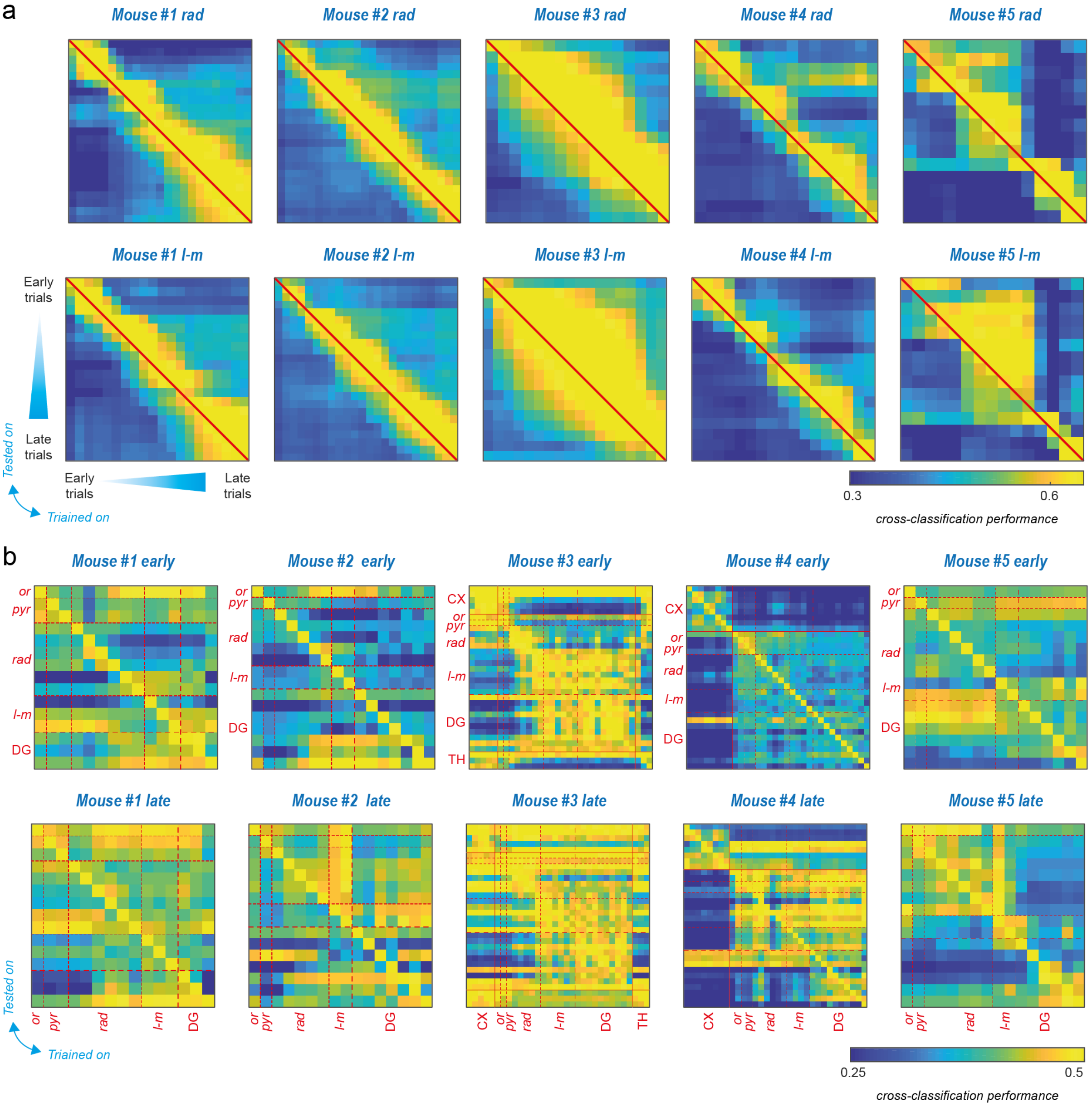


**Extended data Figure 12. Additional details about cross-classification analyses.** Cross-classification matrices for all mice between different trial ranges along learning (**a**, for representative *rad* and *l-m* channels) or between different anatomical locations (**b,** separated for early and late trials). **a,** As in Fig. 4c, cross-classification matrices for classifiers trained on a trial range and tested on another display a characteristic asymmetry between the upper and lower triangular parts, denoting the existence of an “arrow of time” (classifiers trained on future trials can decode information from gamma elements in past trials, but less efficiently the other way around). **b,** As in Figure 4e, cross-classification matrices for classifiers trained on a channel and tested on other channels have a block structure related to anatomical subdivisions. Furthermore, the comparison between cross-classification matrices for early and late trials showed increased cross-classification between blocks (apart from a counter-tendency mouse #5, orange color dot in Fig. 4f).
